# Coupling the role of lipids to the conformational dynamics of the ABC transporter P-gp

**DOI:** 10.1101/2024.04.18.590131

**Authors:** Dario De Vecchis, Lars V. Schäfer

**Affiliations:** Center for Theoretical Chemistry, Ruhr Universitty Bochum, D-44780 Bochum, Germany

## Abstract

The ATP-binding cassette (ABC) transporter P-glycoprotein (P-gp) is a multidrug efflux pump that is overexpressed in a variety of cancers and associated with the drug resistance phenomenon. P-gp structures were previously determined in detergent and in nanodiscs, in which different transmembrane helix conformations were found, “straight” and “kinked”, respectively, indicating a possible role of the lipid environment on the P-gp structural ensemble. Here, we investigate the dynamic conformational ensembles and protein-lipid interactions of the two human P-gp inward-open conformers (straight and kinked) employing all-atom molecular dynamics simulations in asymmetric multicomponent lipid bilayers that mimic the highly specialized hepatocyte membrane in which P-gp is expressed. The two conformers are found to differ in terms of the accessibility of the substrate cavity. The MD simulations show how cholesterol and different lipid species wedge, snorkel, and partially enter within the cavity of the straight P-gp conformer solved in detergent. However, the access to the cavity of kinked P-gp conformer solved in nanodiscs is restricted. Furthermore, the volume and dynamic fluctuations of the substrate cavity largely differ between the two P-gp structures, and are modulated by the presence (or absence) of cholesterol in the membrane and/or of ATP. From the mechanistic perspective, our findings indicate that the straight conformer likely precedes the kinked conformer in the functional working cycle of P-pg, with the latter conformation representing a post substrate-bound state. The inaccessibility of the main transmembrane cavity in the kinked conformer might be crucial in preventing substrate disengagement and transport withdrawal. Remarkably, in our unbiased MD simulations, one transmembrane portal helix (TM10) of the straight conformer underwent a spontaneous conformational transition to a kinked conformation, underlining the relevance of both conformations in a native phospholipid environment and revealing structural descriptors defining the transition between two P-gp conformers.

## Introduction

ATP-binding cassette (ABC) transporters constitute a superfamily of proteins essential for the translocation of a wide range of compounds across cellular membranes, powered by the hydrolysis of ATP.^1–4^ Among these transporters, permeability glycoprotein (P-gp), also known as ABC sub-family B member 1 (ABCB1) or multidrug resistance protein 1 (MDR1) is a pivotal multidrug efflux pump known for its role in drug resistance, particularly in various cancers where it is frequently overexpressed.^5^ Thus, understanding the structural dynamics and functional regulation of P-gp is desirable for understanding the mechanisms underlying MDR and for devising strategies to possibly overcome it.

P-gp exercises its prime role in cellular detoxification,^6^ but it has been proposed to also operate as a low-rate lipid floppase (outward-directed flippase).^7–9^ Hence, phospholipids could effectively play a dual role in P-gp regulation, serving as transport substrates and providing the molecular membrane milieu for the externalized compounds, as suggested by NMR ^10^ and molecular dynamics (MD) simulations.^11,12^ In line with this, the lipid composition of the membrane has emerged as a critical determinant of P-gp function,^13^ with alterations in membrane biophysics affecting P-gp function^14^ demonstrating implications for multidrug resistance.^15^

P-gp is expressed in systems which are specifically devoted to create a functional barrier and detoxify the organism such as intestinal epithelium, liver hepatocyte, kidney and the blood-brain barrier.^16^ The membranes of these barriers are highly specialized and often rich in cholesterol and sphingolipids.^17^ In the kidney, P-gp is expressed in the brush border membrane, which has an increased cholesterol-to-phospholipid and sphingomyelin-to-phosphatidylcholine ratio compared to the basolateral membrane.^18^ In the liver, the small canalicular (apical) membrane of hepatocytes form a luminal meshwork of tubules between adjacent hepatocytes and is the site of primary bile formation.^19^ The canalicular membrane is harnessed with specialized ABC transporters^20^ and defined by microdomains with high cholesterol and sphingolipid content^21,22^ that could be critical as a mean to withstand the harsh biliary molecular soap.

Cholesterol and sphingomyelin are two membrane components that were shown to determine substrate availability and govern P-gp functional localization.^23–25^ Several studies have been carried out to clarify the still not entirely resolved role of cholesterol,^26^ also recently using MD simulations.^27^ According the *cholesterol-filling model*, the presence of cholesterol in the substrate binding site would play a role in stimulating the P-gp substrate-dependent ATPase activity for small drugs.^28^ Finally, P-gp substrate binding and transport are sensitive to cholesterol depletion/replation (addition of cholesterol to previously cholesterol-depleted system),^29^ as well as its ATPase activity, which increases as the concentration of cholesterol increases and peaks at 30%.^30^

Sphingolipids are also essential substrates and modulators of P-gp activity, impacting the protein drug efflux mechanisms and cellular responses to diverse substrates.^9,31,32^ Moreover, the sphingolipid composition of several MDR cell lines is usually severely altered from that of normal cells.^33^ A work investigated how targeting sphingolipid cellular signaling can effectively alter the basal P-gp activity and improve delivery of small-molecules to the brain.^34^ Sphingolipid analogues were also found to modulate P-gp activity and reduced resistance to the chemotherapeutic agent doxorubicin,^32^ highlighting the functional link between P-gp–lipid interactions and drug efflux.

From a structural biology perspective, the community has made tremendous progress, through the determination of numerous high-resolution P-gp structures in both inward-open^35–43^ and outward-open conformations.^36,43,44^ The majority of these structures have been derived from mouse and other organisms,^43,45^ employing lipid-detergent mixtures that, although providing an environment that is a quite different from a cellular membrane, can provide a valuable means to study ABC transporters *in vitro* and *in vivo*.^46,47^ The recent advances in cryo-electron microscopy (cryo-EM) combined with nanodisc technology, have facilitated the acquisition of high-resolution structural data of P-gp in a defined lipid environment.^48–50^ However, at the stage of our research the complex *in vivo* composition discussed above, including the asymmetric lipid composition in the two leaflets, has not been implemented in nanodiscs. Discrepancies persist between P-gp structures solved in detergent and in nanodiscs, especially concerning variations in the inward-open conformation of the transmembrane portal helices TM4 and TM10. Apart from few exceptions, such as the ABCB1-related transporter Atm1 from *Saccharomyces cerevisiae*,^51^ structures of P-gp solved in nanodiscs consistently exhibit kinked helices TM4 and TM10. In contrast, P-gp structures solved in detergent mixtures predominantly show straight helices, albeit with a few exceptions. Ligand-binding induced kinking of TM4 was previously reported for mouse P-gp^42^ and for the TM10 of the ABCB1-related transporter from *Caenorhabditis elegans*,^45^ both in detergent.

Finally, structural data from hybrid human/murine P-gp constructs indicate conformational flexibility in TM4 and TM10 and an equilibrium between straight and kinked conformations,^35^ and also highlight the significance of the membrane environment for this structural divergence in modulating functional dynamics of P-gp. Studying such dynamic structural phenomena requires very high resolution in both space and time, which can not be fully achieved by current experimental techniques alone. Therefore, open questions remain concerning the precise mechanisms that govern the conformational ensemble and transitions, the effects of the lipid environment, and their functional implications.

MD simulations can answer these open questions by providing the missing atomic-level insights into the structural dynamics of the proteins in their (close to) native lipid environment.^52,53^ Specifically for ABC transporters, simulations have contributed to an increased understanding of their – inherently dynamic – functional working cycles.^54–56^ In some cases, MD simulations could even describe the large-scale alternating access conformational transitions between inward-and outward-facing conformations and their coupling to distinct substrate transport events, such as, for example, the unbiased multimicrosecond simulations of the heterodimeric ABC exporter TM287/288.^57,58^ For P-pg, computational investigations have provided insights into the conformational flexibility of the membrane-embedded transporter, offering valuable insights into its structural dynamics, accessibility of its transmembrane cavity to lipids and other membrane components, and substrate binding.^11,12,27,36,59–66^ Here we employ MD simulations to investigate P-gp structural dynamics in asymmetric multi-component bilayers that mimic the canalicular membrane of hepatocytes. The structural determinants governing the two distinct P-gp inward-open conformations, straight and kinked, are compared. The simulations capture conformer-specific preferential P-gp– phospholipid and P-gp–cholesterol interactions, shedding light on the differential interactions with the membrane components. Notably, several residues that are found to be preferential lipid interaction sites are linked to cancerous mutations. Phospholipid and cholesterol molecules enter the P-gp main transmembrane cavity of the straight but not of the kinked conformer, suggesting a blockage to the main substrate cavity in the kinked conformation. Our results highlight the role of the membrane environment in modulating the conformational ensemble and flexibility of P-gp, in particular, of the helices TM4 and TM10. We record a conformational change of TM10 from straight to kinked and pinpoint conserved residues that might act as structural tipping points. Finally, a mechanistic scheme is proposed for the functional role of the two inward-open conformers, straight and kinked, providing insights into the transport cycle and reconciling available experimental data.

## Methods

### Molecular Modeling

The coordinates of the nanodisc-reconstituted drug-free human P-gp (PDB 7A65,^48^ resolution 3.9 Å) were loaded in ChimeraX and ISOLDE^67^ was used in combination with the associated cryo-EM density map EMD-11666 to inspect the density and perform local refinement of the structure. Then, the MolProbity server^68^ was used to assess the structure refinement procedure that finally led to a MolProbity score of 1.17 (initial score was 2.06), a clash score of 0.6 (initial was 13.76), a decreased number of poor side chain rotamers (from initial 16 to final 11), an increased number of favored rotamers (from initial 883 to final 895) and no Ramachandran outliers. This structure, in combination with residues 77–110 from the human P-gp cryo-EM structure solved in the presence of the drug paclitaxel (PDB 6QEX,^50^ resolution 3.6 Å), was used to build the final apo-P-gp structure and to revert the mutation A893S present in the structure 7A65 to the wild-type sequence using 500 iterations with Modeller.^69^ This was done to reconstruct the external transmembrane (TM) helices TM1 and TM2 (C- and N-terminal, respectively), not present in PDB 7A65. Similarly, 500 iterations of Modeller were performed to build holo-P-gp (i.e., in complex with Mg-ATP). The ATP and Mg coordinates and the interacting residues 401–409, 427–435, 474–476, 555–557, 1043–1052, 1070–1078, 1117–1119 and 1200–1201 were taken from the cryo-EM structure of the ATP-bound outward-facing conformation (PDB 6C0V,^44^ resolution 3.4 Å). The procedure described above generated apo- and holo-P-gp models with the portal helices TM4 and TM10 in the kinked conformation. These models were further processed to also build apo- and holo-P-gp models with the portal helices in the straight conformation. Firstly, the Cα atoms of residues 204–270 (TM4 and ICH2) and 851–914 (TM10 and ICH4) from the X-ray crystal structure of murine P-gp (PDB 4M1M,^70^ resolution 3.8 Å) were aligned to the Cα atoms of residues 208–274 (TM4) and 855–918 (TM10) of human P-gp, respectively. Secondly, only residues 208–255 and 851–897 from 4M1M were used as template to model the TM4 and TM10 helices of human P-gp, that is, the ICHs where only considered to obtain a better structural superposition. Finally, the obtained patched straight structures were subsequently refined with ISOLDE in combination with the cryo-EM density map from human P-gp (EMD-4284, resolution 4.5 Å) that has straight TM4 and TM10 helices.^35^ During the refinement with ISOLDE, all coordinates but TM4 and TM10 helices were restrained. Therefore, apart from the TM4 and TM10 coordinates, the kinked and straight P-gp models are very similar.

### Molecular dynamics simulations

The sizes and compositions of all the simulated systems are listed in Table 1. All simulations were carried out with GROMACS version 2021.1.^71^ The P-gp structural models obtained as described above were initially converted to the coarse-grained (CG) Martini (version 2.2) force field^72^ using the *martinize.py* script. At this stage, the ATP and Mg coordinates were not considered. The tool *insane* was used to generate three independent membrane assemblies and insert the protein into an asymmetric bilayer resembling the composition of the bile canaliculi plasma membrane from the hepatocyte.^22^ The extracellular leaflet is composed of 40% cholesterol (CHOL), 27% 1-palmitoyl-2-oleyl-phosphatidylcholine (POPC), 23% sphingomyelin (SM) and 10% 1-palmitoyl-2-oleyl-phosphatidylethanolamine (POPE). The intracellular leaflet is composed by 40% cholesterol, 17% POPE, 15% 1-palmitoyl-2-oleyl-phosphatidylserine (POPS), 10% phosphatidylinositol bisphosphate (PIP_2_), 8% POPC and 10% SM. Each leaflet has ∼200 lipids (see Table S1). Similarly, the protein was also embedded in a membrane deprived of cholesterol. In these systems, the cholesterol molecules were replaced by POPC and POPE phospholipids in the extracellular and intracellular leaflet, respectively. All the systems were neutralized with a 150 mM concentration of NaCl and subsequently energy-minimized with steepest descent (about 1400 steps). After minimization one equilibration of 5 µs was performed for each system using different random seeds for the initial velocities and with harmonic position restraints on all the protein beads (force constants of 2000 kJ mol^−1^ nm^−2^). The time step used for integrating the equations of motion in the coarse-grained simulations was 20 fs. The “new-RF” simulation parameters suggested by de Jong et al.^72^ were used. Equilibrations were performed at 310 K with protein, membrane and solvent separately coupled to an external bath using the v-rescale thermostat with a stochastic term^73^ with coupling time constant τ*_T_* = 1 ps. Pressure was maintained at 1 bar using the stochastic cell rescaling (c-rescale) barostat^74^ with semi-isotropic conditions (coupling time constant τ*_P_* = 12 ps and compressibility 3 × 10^−4^ bar^−1^). Coordinates were saved to the disk every 400 ps. The obtained systems from each conformer (*kinked* and *straight*) were energy minimized (about 600 steps) and then back-mapped using the *backward* method^75^ to all-atom resolution with the Charmm36m all-atom force-field.^76^ Subsequently, the refined apo- and holo-P-gp models generated as explained above, were superposed over the Cα atoms to the back-mapped protein coordinates, and the latter deleted. Thus, the CG simulations were only used to equilibrate the environment of the protein, but the protein coordinates from the CG simulations were not used downstream in the project. Each atomistic system was energy-minimized (steepest descent, about 3000 steps). The parameters for the Mg^2+^ ion were taken from Allńer and colleagues.^77^ Harmonic bonds with force constants of 1000 kJ mol^−1^ nm^−2^ were introduced between the Mg ions and the coordinating Oδ-atoms from the Asn475 and Asn1118 side chains in the NBD domains. The systems were equilibrated for 50 ns with harmonic position restraints on all protein heavy atoms, followed by 100 ns with only the protein backbone atoms restrained (using restraining force constants of 1000 kJ mol^−1^ nm^−2^ in both cases). The equilibrations were performed under NPT conditions at 310 K and 1 bar, using the v-rescale thermostat^73^ (coupling time constant τ*_T_* = 0.1 ps) and the stochastic cell rescaling (c-rescale) barostat^74^ (in semi-isotropic mode, with coupling time constant τ*_P_* = 2 ps and compressibility 4.5 × 10^−5^ bar^−1^). Non-bonded Lennard-Jones interactions were smoothly shifted to zero at a cut-off of 1 nm, which was also used for the real-space part of the Coulomb (electrostatic) interactions. Long-range electrostatics were treated with the particle-mesh Ewald method^78^ with a 0.12 nm grid spacing. The LINCS algorithm^79,80^ was used to constrain bond lengths involving H atoms, allowing an integration time step of 2 fs. For each system a 1 ns pre-run (unrestrained) with 1 fs time step was performed, followed by three independent unrestrained production simulations of 1 µs each using different random seeds for the initial atomic velocities. Parameters and conditions were the same as described for the equilibrations above. Coordinates were saved to disk every 20 fs.

**Table 1:**
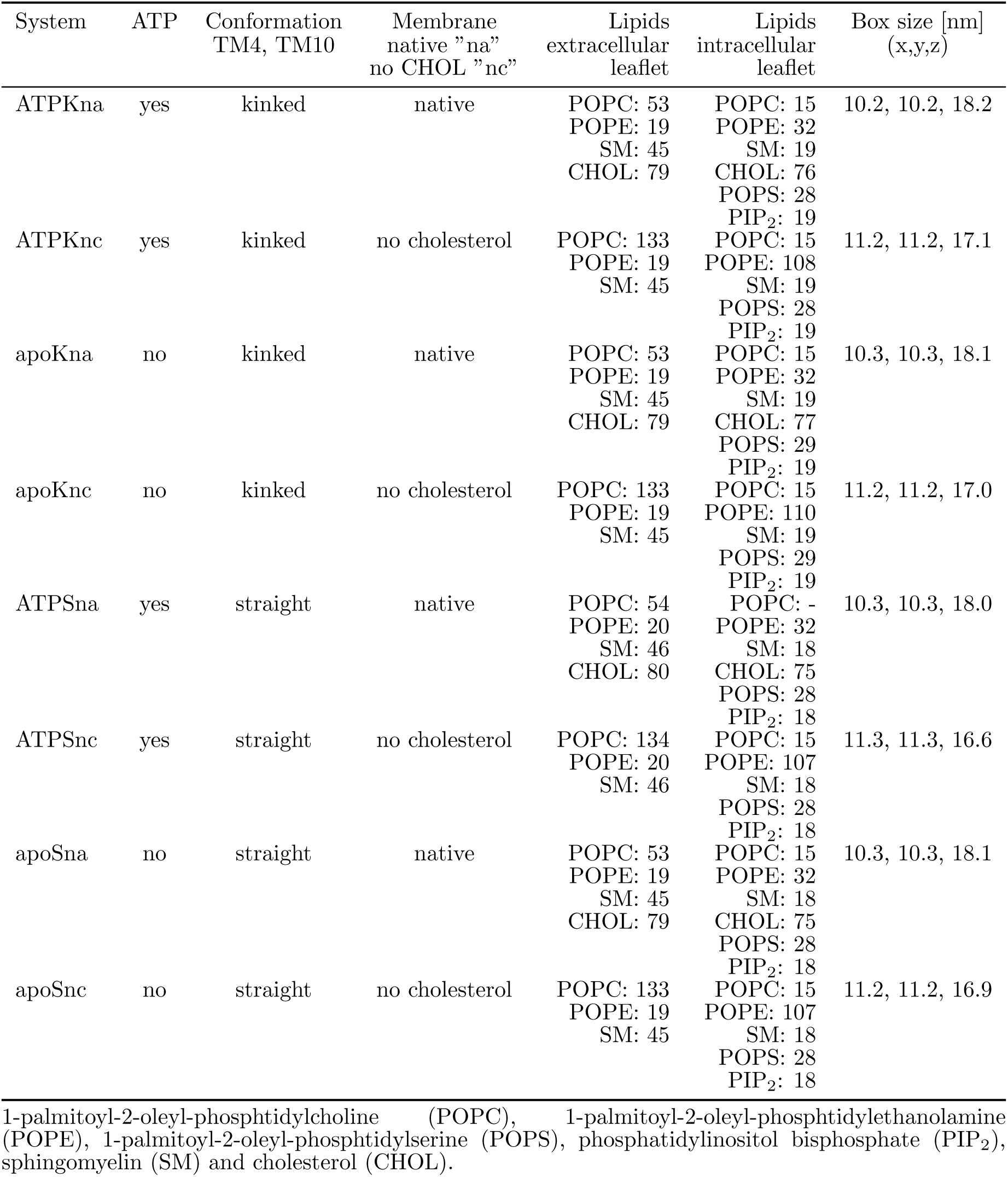
Summary of simulated systems.

### Analyses

For the CG MD simulations, the GROMACS analysis tool gmx densmap was used to calculate the 2D number-density map for each membrane component. For the all-atom MD simulations, the average number of protein-lipid contacts across all the repeats and for each simulated system were calculated with gmx select within a cut-off of 0.55 nm between all the protein atoms and the headgroup atoms of each lipid species and cholesterol. The tool gmx distance was used to calculate all the distances.

The algorithm T-Coffee^81^ was used for the multiple sequence alignment using *slow pair* as pairwise method. Mutation information related to human diseases and cancers were obtained from UniProtKD^82^ and the catalogue of somatic mutations in cancer (COSMIC) database (https://cancer.sanger.ac.uk).^83^

For the all-atom MD simulations, the tool trj cavity^84^ was used to calculate the volume of the P-gp main transmembrane cavity as well as for the membrane component headgroups that overlap with it. The P-gp residues selected for the calculation of the enclosed cavity are 59–81, 107–133, 188–238, 292–318, 328–352, 718–741, 749–774, 827–881, 938–961 and 971–995. For each system, the three independent repeats were first concatenated and the coordinates saved every 200 ps. The trajectories were then fitted on the Cα atoms of the selected residues before the calculation. The options dim 5 and dim 6 (i.e., the degree of burial) with spacing 1.4 (i.e., the size of the grid voxel in Å) were used for the kinked and straight conformer, respectively.

Secondary structure elements were identified using the do dssp tool implemented in GRO-MACS based on the DSSP method.^85^ The initial 100 ns were removed from each of the three independent repeat simulations. Then, the trajectories were concatenated and the coordinates analyzed every 100 ps. An in-house script was used to calculate the percentage of each secondary structure element.

The angle α between the two P-gp transmembrane bundles (bundle 1: TM1, TM2, TM3, TM6, TM10, TM11; and bundle 2: TM4, TM5, TM7, TM8, TM9, TM12) was calculated as follows: First, gmx distance was used to calculate the x,y,z components of the vectors ⃗v_1_ and ⃗v_2_ defined between the center of mass of the Cα atoms of residues 89, 90, 91, 92, 96, 97, 98, 99, 207, 208, 209, 210, 212, 213, 214, 215, 320, 321, 322, 323, 330, 331, 332, 333, 737, 738, 739, 740, 745, 746, 747, 748, 850, 851, 852, 853, 855, 856, 857, 858, 962, 963, 964, 965, 971, 972, 973, 974 (see Fig. S1e, residues shown in blue) and the center of mass of the Cα atoms of residues 154, 155, 156, 157, 168, 169, 170, 171, 367, 368, 369, 370, 899, 900, 901, 902, 913, 914, 915, 916 (see Fig. S5b, residues shown in orange) and 255, 256, 257, 258, 270, 271, 272, 273, 795, 796, 797, 798, 811, 812, 813, 814, 1010, 1011, 1012, 1013 (see Fig. S1e, residues shown in green), respectively. Second, the angle α between these two vectors was calculated, 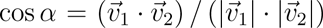, where 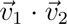 ⃗is the scalar product and 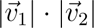 is the product of the norms.

## Results

### Phospholipids and cholesterol accumulate at P-gp portals in both kinked and straight conformers

First, we aim to elucidate the conformational dynamics of both the straight and kinked P-gp conformers, to investigate whether, and if so how, the lipids that constitute the highly specialized hepatocyte bile canaliculi membrane selectively interact with these inward-open conformers. The kinked and straight P-gp conformers are shown in Fig. 1a,b.

**Figure 1:**
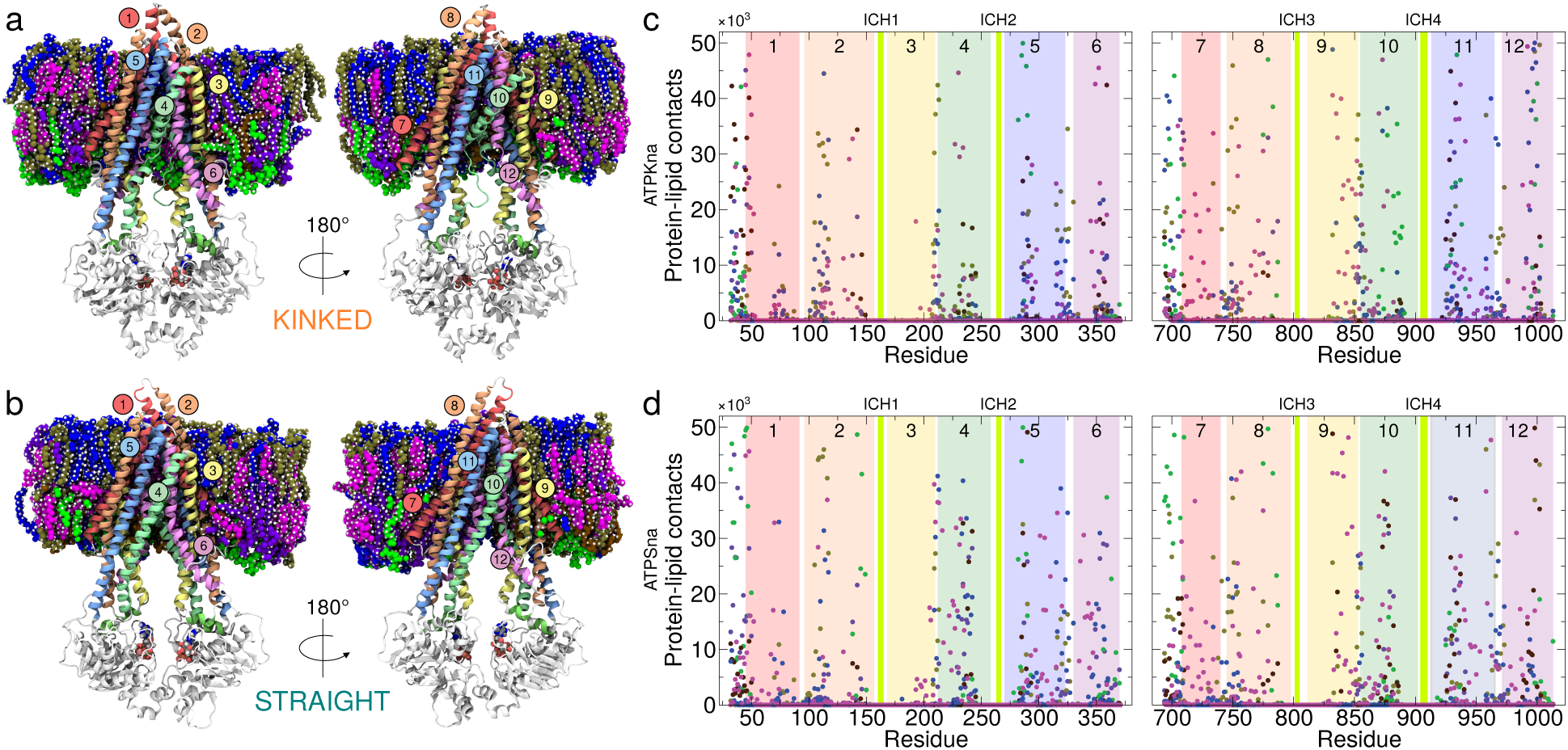
Membrane-embedded P-gp in the two different inward-open conformations. (a) Simulation snapshot of the P-gp kinked conformer embedded in the hepathocyte plasma membrane model. The snapshot is from the ATPKna system (see Table 1). The protein is shown in ribbon with the transmembrane helices numbered and colored in rainbow. The ATP and the membrane components are shown in van der Waals representation, cholesterol in magenta, sphingomyelin in blue, phosphatidylcholine in tan, phosphatidylethanolamine in violet, phosphatidylserine in brown, phosphatidylinositol in green. Solvent and ions are not shown for clarity. (b) As in a but for the P-gp straight conformer. The snapshot is from the ATPSna system (see Table 1). (c) Protein-lipid contact analysis of the P-gp kinked conformer. Phospholipid-P-gp and cholesterol-P-gp contacts during the accumulated 3 µs of all-atom MD simulation time of the ATPKna system for TM1 to TM6 (residues 22–381) and TM7 to TM12 (residues 684–1023). The P-gp transmembrane helices are numbered and colored as in a and b. The color legend for the lipids and cholesterol is the same as in a and b. Intracellular coupling helices ICH1 to ICH4 are indicated in green. (d) As in c but for the P-gp straight conformer (ATPSna system).

For both conformers, during the initial coarse-grained MD equilibrations, phospholipids from the lower leaflet were observed to “snorkel” at the portals, that is, the lipid headgroups partially enter and exit the transmembrane cavity of P-gp in a dynamic manner, but the lipids did not fully enter the cavity and bind inside it (Fig. S2). Then, the coarse-grained simulation systems were back-mapped to all-atom resolution and the protein coordinates were exchanged with the apo- and holo-P-gp models generated initially from the experimental structures, to remove any bias from the coarse-grained model on the protein coordinates (see Methods). The subsequent all-atom MD simulations reveal a complex network of distinct protein-lipid contacts (”fingerprints”^86^) for both conformations (Fig. 1c,d).

Globally and among all the simulated systems, for POPC most of the protein-lipid interactions involve the portion of the transmembrane helices exposed to the extracellular side of the membrane, with additional contribution from the cytoplasmic-facing C-term elbow helix (residues 694–707), TM5, TM9, TM11 and TM12 (Table S1 and Fig. S3). Similarly, for POPE, most interactions map on the cytoplasmic side among the N-term (residues 32–44) and C-term elbow helices, TM1, TM4, TM5, TM6, TM7, TM8, TM9, TM10 and TM11 (Table S1 and Fig. S3). For POPS, interactions were found across all the TM helices, including residues at the N-term and C-term elbow helices (Table S1 and Fig. S3). Interactions with PIP_2_ were found across all the TM helices, including both the N- and C-term elbow helices, with the exception of TM3 and TM9, which remain rather shielded (Table S1 and Fig. S3). As expected from the higher concentration of SM in the extracellular leaflet of the bilayer, P-gp interactions with SM are rather distributed across all the extracellular portions of the TM helices. Nevertheless, contacts above 60% threshold were also found at the cytoplasmic side of both P-gp lateral portals and involving TM4, TM6, TM9, TM10 and TM12. In addition, in the simulations of the cholesterol-free membrane, an overall reduction in the number of residues interacting with SM was observed (Table S1 and Fig. S3). Finally, the presence of ATP does not seem to affect the P-gp–cholesterol interactions, which remain distributed across all the TM helices, even though some preferential contact residues were identified (Table S1 and Fig. S3).

We further expanded our investigation and found 17 P-gp residues in the transmembrane domain that preferentially interact with lipids. Interestingly, mutations of these residues were also found to be associated with various types of cancers (Table S1 and Fig. S4). Similarly, the MD simulations pinpoint 11 residues among natural variants, involved in an altered drug response to tramadol, a common analgesic. Of notice, mutants were also found on the portal helices TM4 (A230V), TM6 (A356S), TM10 (V873G, V874I, M876R) and TM12 (A999T), see Table S1 and Fig. S4.

The analysis of the protein-lipid contacts reveals distinct groups of P-gp residues that are involved in membrane interactions. These groups of residues do not overlap between the two inward-facing conformers, kinked and straight, and thus represent a conformer-specific “fingerprint”. Different non-overlapping groups of lipid-contacting P-gp residues were also identified in absence of ATP or when no cholesterol was present in the membrane. Moreover, our results suggest a role in lipid transport, and possibly cancer development, for residues located at the lateral portals of the P-gp transmembrane domain.

### Conformational change via structural tipping points in helices 4 and 10

Next, we turn to the P-gp main transmembrane cavity, which is enclosed by TM4 and TM10 together with TM6 and TM12 (Fig. 2a, b). Structural data suggest a concerted movement of these helices to enclose different P-gp transport substrates and inhibitors such as zosuquidar and tariquidar.^35,42,48,87^ Therefore, it is important to compare the dynamics of this volume between the conformers and to map the available druggable space. Under native conditions, that is in presence of ATP and cholesterol, for the kinked conformer the volume is in the range 1000–2500 Å^3^ (Fig. 2a, b orange lines), that is, it dynamically fluctuates around the initial value of ∼1700 Å^3^ (Fig. 2a, b dotted orange vertical line).

**Figure 2:**
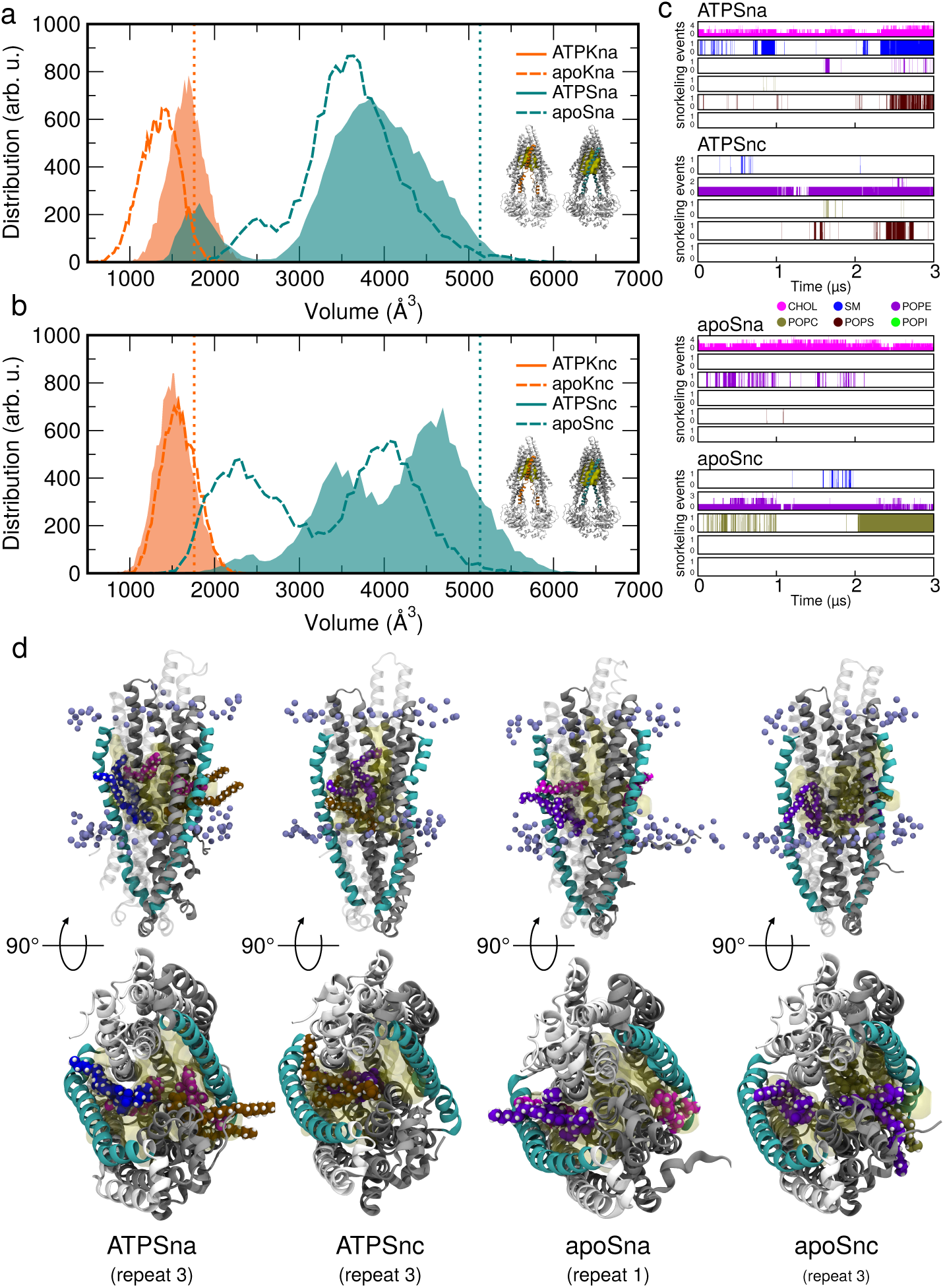
Characterization of the P-gp main transmembrane cavity. (a and b) Distribution of the volume of the P-gp transmembrane cavity during the MD simulations. The initial volume is indicated by vertical dotted lines. The insets on the right are the initial P-gp conformer models, the kinked conformer on the left (orange) and the straight conformer on the right (cyan). The volume of their respective cavity is shown in yellow and both TM4 and TM10 are indicated. (c) Cholesterol and phospholipid snorkeling events from the membrane into the P-gp cavity during the MD simulations. A snorkeling event was recorded if the phosphorous or the oxygen atoms from phospholipids or cholesterol, respectively, were positioned inside the cavity volume shown in a, b. The three repeat simulations (of 1 µs each) were concatenated. The color legend is the same as in Fig. 1. (d) Snapshots from simulations showing the transmembrane cavity of the straight conformer with the membrane components (in van der Waals representation), snorkeling within the cavity (indicated in yellow). The phosphorous atoms from the lipid headgroups are indicated as spheres. Solvent, ions and NBD domains have been removed for clarity.

For the straight conformer the volume of the cavity is much larger and deviates from the initial value of ∼5100 Å^3^ (Fig. 2a, b dotted cyan vertical line). The straight conformer has a much wider distribution with an average volume of ∼4000 Å^3^, and the cavity volume even reaches values up to 6000 Å^3^ (Fig. 2a, b cyan lines). Interestingly, in the native membrane the cavity volume of both conformers decreases in absence of ATP (Fig. 2a dashed orange and cyan lines), which is somewhat counter-intuitive. For the straight conformer, this effect is confirmed when cholesterol is removed (Fig. 2b, dotted cyan line). The simulations indicate that the different conformations of the helices TM4 and TM10 impact the volume distribution of the P-gp transmembrane cavity, and suggest a relation between this volume and the presence of ATP.

Beyond xenobiotics and drugs, P-gp is known to bind and extrude phospholipids^8,9,88^ and several works have shown that the transmembrane cavity is accessible to different membrane components.^11,60,61^ Therefore, to further investigate whether the two conformers are permeable and might differ in their lipid accessibility profiles, the number of cholesterol and phospholipid molecules that snorkel inside the P-gp cavity were analyzed (Fig. 2c, d). A snorkeling event was counted if the phosphorous atom or the oxygen atom of the phospholipid headgroups or cholesterol, respectively, were found to be positioned inside the cavity volume. For the kinked conformer, even though lipids and cholesterol accumulate at both TMD portals (Fig. 1c, d), the cavity remains sealed and no snorkeling events were detected. Conversely, during all the simulations of the straight conformer, with the exception of PIP_2_, phospholipids and cholesterol were observed at both portals between TM4/TM6 and TM10/TM12 and snorkel with their headgroups into the main P-gp cavity (Fig. 2c, d). This result suggests that, for the initial loading of the transmembrane cavity with substrate, P-gp needs to be in the straight conformation (see discussion below).

Remarkably, in the second repeat simulation of the apoSnc system (after ∼170 ns), TM10 underwent a structural transition from a straight to a kinked conformation (Fig. S5a, Video S1). The RMSD of the TM10 Cα atoms with respect to the initial P-gp kinked structural model and the ATPKna simulated system is 3.7 Å and 2.6 Å, respectively (Fig. S5). Following this conformational change, and in line with the previous results, the TM10/TM12 portal becomes sealed to the lipids which otherwise were observed to enter the cavity in the other repeats (Fig. 2c, d and Fig. S5b). The structural change detected in this “half-straight/half-kinked” conformer reduces the cavity volume, explaining the lower values observed for the apoSnc system (Fig. 2b, dotted cyan line). Interestingly, after the straight-to-kinked conformational transition of TM10, in the simulation the phospholipids are still able to enter the transmembrane cavity from the opposite portal formed by TM4/TM6 (Fig. S5b).

To more systematically analyze the integrity of the P-gp helices during the simulations, we analyzed the secondary structure of TM4 and TM10 for all the simulated systems (Fig. 3). These analyses pinpoint the residues that are the tipping points behind the observed conformational change. TM10 appears to be the most flexible in both conformers. In particular, for the kinked conformer a stretch of mostly coil residues interrupts the helix at position 880–890, and at position 869–872 and 877–892 for the straight conformer. The apoSna system shows in this region a somewhat frustrated TM10 helix, in particular around residues 877–892 (Fig. 3). Although the conformational change was only observed in the apoSnc system, residues 869–873 in TM10 are partially unwinding in almost all the straight P-gp systems, as is also the case for residues 863–864, the latter showing a similar behavior as in the kinked conformer (Fig. 3). Modest helix distortion/unwinding was also detected for TM4 around residues 220–222 and residues 246–247, especially for the kinked conformer and the apoSnc system (Fig. 3).

**Figure 3:**
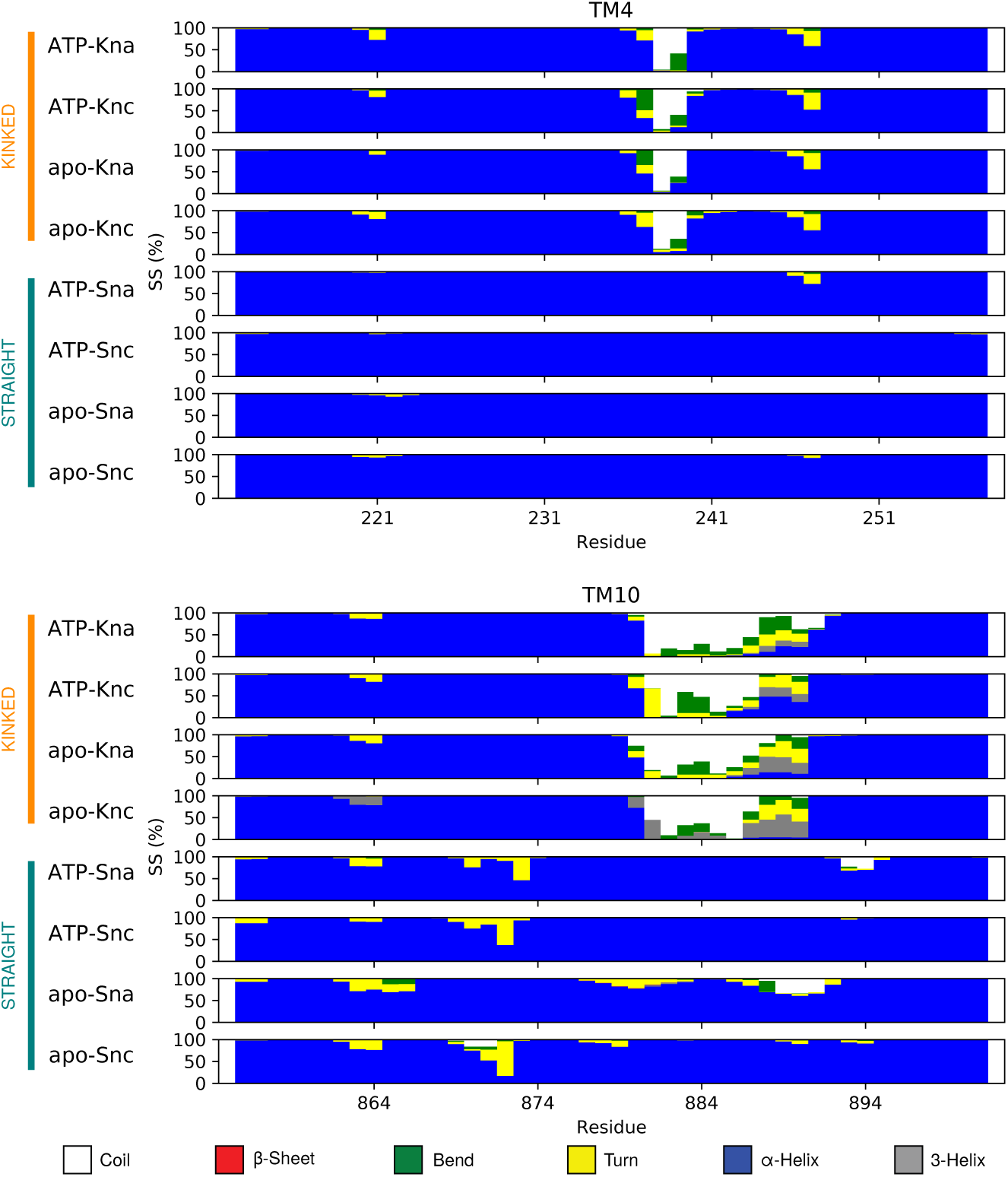
Evolution of the secondary structure of TM4 and TM10 in the simulated systems. The plots show the persistence of each secondary structure element (color code at the bottom) calculated as a percentage during all concatenated repeat simulations.

In summary, even though both conformers are in an inward-open conformation our results indicate that the lipid accessibility of the main cavity differs. Larger portals are suggested for the P-gp straight conformer, and bending of helices TM4 and TM10 can govern the accessibility of the transmembrane cavity. Finally, a partial transition was observed from the P-gp straight to the kinked conformer via tipping points located within helices TM4 and especially TM10, possibly hinting at a role of the membrane environment in modulating the P-gp conformational ensemble.

### Correlation between opening of transmembrane bundle and substrate accessibility of main cavity

Our previous findings demonstrate that the kinked conformation limits access to the main cavity. To study P-gp flexibility various descriptors have been used, such as the distance between the NBD domains^89^ and the angle between the TM helix bundles.^66^ We aim to investigate whether these descriptors differ between the straight and kinked inward-open conformers, and how they correlate with the transmembrane cavity volume, which differs between the two conformers (Fig. 2). Therefore, we further analyzed how the accessibility of the transmembrane cavity is coupled to the global conformational dynamics of the transporter.

For each simulated system, the angle α between the two P-gp transmembrane bundles (Fig. S1) and the distance between the center of mass (COM) of the two NBD domains (Fig. S6) was calculated, see Methods. The results show comparable values for both conformers when simulated in a native membrane and in complex with ATP, with an angle of ∼32^◦^ (Fig. S1a) and an NBD dimer distance of ∼3.5 nm (Fig. S6a). These average values are the same in the simulations without cholesterol (Figs. S1b, S6b). As expected, the absence of ATP loosens the NBD dimer and increases protein flexibility. This effect might be slightly more pronounced for the kinked conformer, for which the NBD dimer distance transiently fluctuates up to values of ∼5 nm (Fig. S6c). Interestingly, as shown above, the volume of the main cavity is decreased in the apoKna system (Fig. 2a, orange dashed line), questioning whether the NBD dimer distance is a good reporter (or proxy) for transmembrane cavity accessibility (see below). In the absence of ATP and cholesterol, an increase in protein flexibility is detected for both conformers, with a transient maximum NBD dimer distance of ∼5.8 nm and the angle α of ∼40^◦^ for the kinked (Figs. S1d, S6d repeat 1, after 100 ns) and ∼5.5 nm and ∼37^◦^ for the straight conformer (Figs. S1d, S6d repeat 2, after 50 ns).

We further analyzed the correlation between the NBD dimer distance and the opening angle of the P-gp transmembrane bundles (Fig. S7). In general, there is a moderately positive correlation between the two descriptors. Thus, as expected, an increase of the NBD-NBD distance is correlated to a larger opening angle α.

Remarkably, in the first repeat simulation of the apoSna system, a negative correlation coefficient of -0.34 is found (Fig. S7a). Close inspection of the trajectory revealed that in this simulation, the NBD2 detached from both ICH2 and ICH3. As a consequence, the transmembrane bundle collapsed somewhat (low α values), and uncouples from the NBDs. In particular, the two charged residues Arg262 and Asp805 (from ICH2 and ICH3, respectively) loose contacts, ultimately leading to the detachment of NBD2 from ICH2/3. We interpret this as a simulation artefact (probably linked to force field imperfections^90^), which however confirms the functional importance of the ICHs as conformational transducers between the NBDs and the transmembrane bundles.

We finally investigated how the transmembrane cavity volume, which differs between the two conformers, correlates with the opening angle α (Fig. 4) and the NBD-NBD distance (Fig. S7b). For the kinked conformer the analysis indicates that despite the fluctuations observed for the two descriptors used (see above), the cavity volume remains tight, with no (or even slightly negative) correlation between the two variables for the system under native condition, ATPKna (Fig. 4 and Fig. S7b). For the straight conformer the distributions of the cavity volume are broad (Fig. 2), and overall positive correlation is found, especially for the ATPSnc system (Fig. 4 and Fig. S7b).

**Figure 4:**
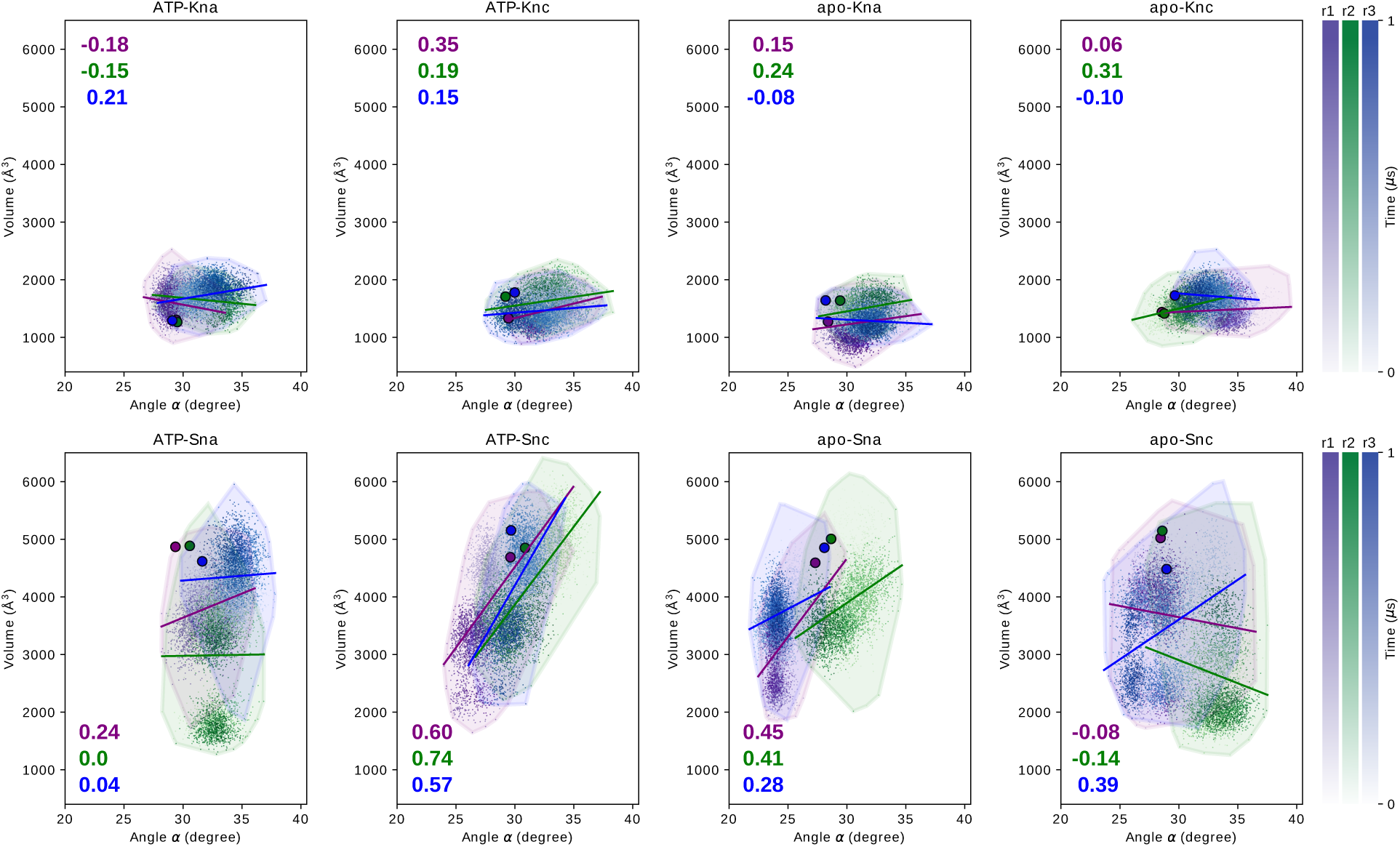
Correlation between the P-gp transmembrane cavity volume and the angle α between the transmembrane helix bundles. For each simulated system and each simulation repeat, the Pearson correlation coefficients are indicated. The three simulation repeats are shown in different colors, with the color gradient indicating the simulation time. The circles indicate the initial values at the beginning of the simulations.

The results reported above indicate that the P-gp transmembrane domain is flexible and can reach wider degrees of opening (with respect to the starting structures of the simulations). This is also true for the kinked conformer that, although flexible, retains TM4 and TM10 in kinked conformations (Fig. 3) and the main cavity sealed from the membrane environment. Regarding the 2nd repeat simulation of the apoSnc system at the time of the conformational change of TM10 (i.e., after 170 ns), the angle α is ∼30^◦^ (Fig. S1) and the NBD distance is ∼4 nm (Fig. S6). Since these values are also found among the other simulated systems, and in light of the only moderate correlation of these two structural descriptors with the cavity volume (see above), we further aimed to identify alternative structural descriptors to distinguish the two conformers. We initially considered the Cα–Cα distance between residues Lys147 (TM2) and Met791 (TM8), which have been previously used as spin-label positions in electron paramagnetic resonance (EPR) studies of the conformational flexibility of murine P-gp (Gln145–Met787^91,92^ see Fig. S8). Even though somewhat more broadly distributed for the straight conformer compared to the kinked (Fig. 5a, bottom panel), the values are similar, especially for the native membrane conditions (Fig. 5a, dark green line), and compatible with the values of ∼45 Å observed for mouse P-gp.^91,92^ However, for the TM4–TM10 residue pair G226–G872, which is the pivotal region underneath the kink in the observed straight-to-kinked conformational transition (Figs. 3, 5 and Fig. S5), distinct values for the two conformers were observed. In particular, a sharp distance distribution around 2 nm for the kinked conformer (Fig. 5a, orange lines), and in the range 2.9–3.5 nm for the straight conformer (Fig. 5a, cyan lines). Fig. 5b shows how the backbone at position 872 pivots to enable TM10 bending towards the transmembrane cavity. Analysis of the conservation of residues Gly226 and Gly872 between different P-gp homologous proteins (Fig. S8) reveals a high degree of conservation for both residues among higher eukaryotes. Noticeably, within TM4 and TM10, highly conserved prolines are present at positions -3 and -6 from Gly226 and Gly872, respectively, which might as well act as pivots during the conformational transition as previously reported for the mouse P-gp.^35^

**Figure 5:**
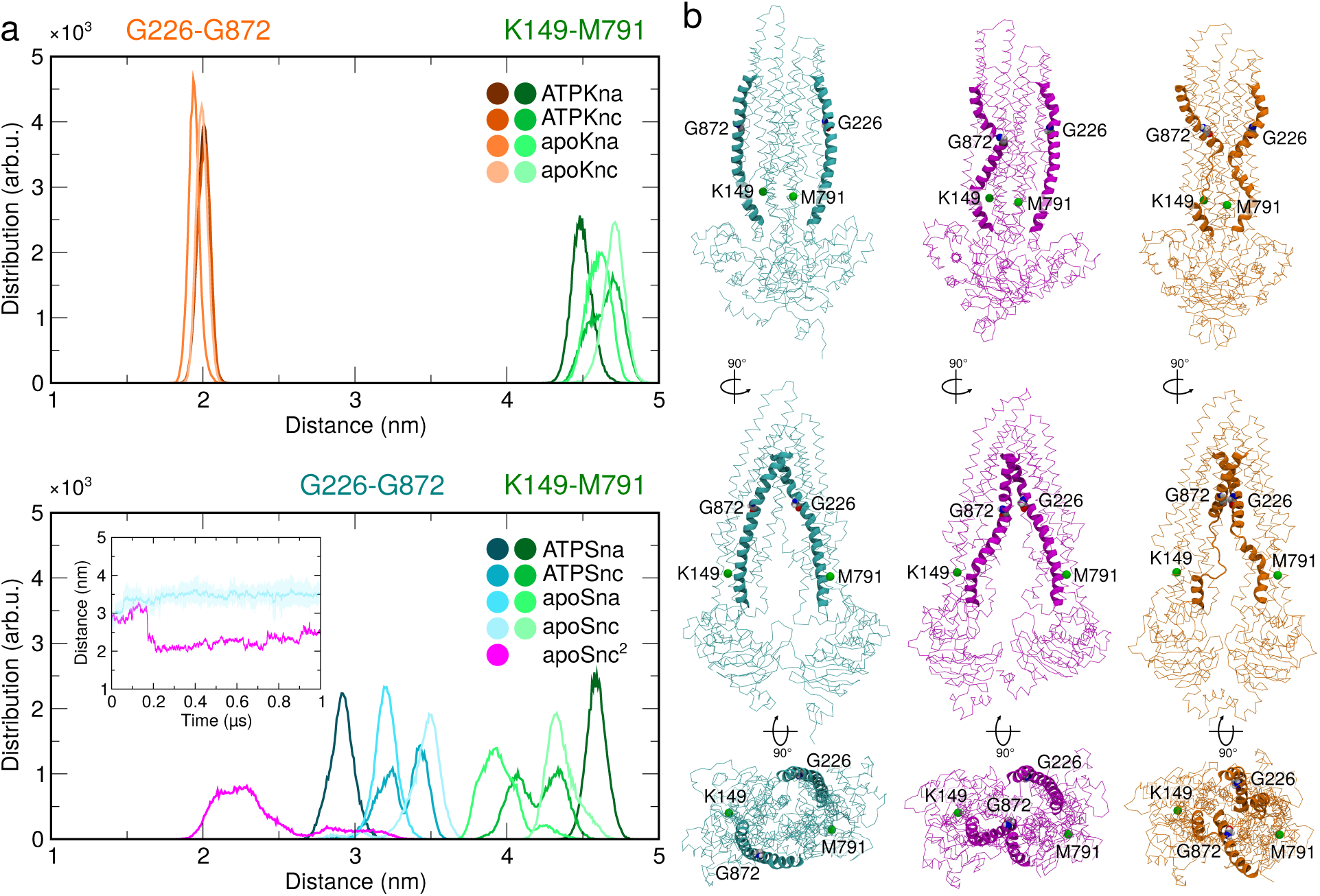
P-gp structural descriptors define alternative conformational states. (a) Distance distributions between residue pairs for the kinked (top) and straight (bottom) conformers. The inset shows the time trace for the “repeat 2” simulation of the apoSnc system in which the half-straight/half-kinked conformational transition occurs (magenta), in comparison to “repeat 1” (cyan). (b) Snapshots from simulations of the straight (cyan), kinked (orange) and half-straight/half-kinked (magenta) conformers. The residues used for the distance calculations are indicated.

In conclusion, our analyses underline that one might have to be cautious with the intuitive assumption that a larger transmembrane bundle angle α and a larger distance between the NBDs indicate increased substrate accessibility of the P-gp transmembrane cavity. Especially for the kinked conformer, this assumption might not be valid. Instead, the distance between residues Gly226 and Gly872 might be a preferable descriptor to distinguish not only the two stable conformers (kinked and straight), but also to describe the transition between them. Therefore, it could be worthwhile to consider these residues as EPR spin label positions.

## Discussion

In this study, all-atom MD simulations are employed to examine the structural dynamics of two inward-open P-gp conformers, kinked and straight, which largely differ in the conformation of their portal helices TM4 and TM10. Apart from these two transmembrane helices, the structures of the two P-gp conformers are similar. The structures were solved under different conditions, detergent for the straight conformer whereas the kinked structure was obtained in nanodiscs. This pattern was recently confirmed across species, as evidenced by the straight conformation observed in the mouse P-gp structure solved in detergent.^36^ In our simulations we mimicked the highly specialized canalicular membrane composition, enriched with sphingomyelin and cholesterol, which can withstand the harsh environment formed by the bile molecular soap.^17^ To better understand the dynamics and structural relationships of helices TM4 and TM10 under various conditions, the two conformers were simulated under native-like conditions and in different environments by removing cholesterol and ATP, known regulators of P-gp dynamics, individually and in combination, to explore their effects on the structural dynamics and conformational ensembles.

### P-gp dynamics in a lipid context

The MD simulations revealed a complex network of protein-lipid interactions that reflects the membrane’s pronounced asymmetry. Phospsholipids and cholesterol accumulated at the P-gp portals exhibiting dynamic “snorkeling” from the intracellular leaflet, in line with previous coarse-grained simulations.^60^ Apart from few exceptions, our MD simulations revealed a unique pattern of protein-lipid contacts for both P-gp conformers. Interactions with different lipids were mapped across various transmembrane helices, with preferences observed for specific regions depending on the lipid type. Anionic phospholipids present in the intracellular leaflet (PIP_2_ and POPS) and cholesterol were found to strongly interact with all P-gp TM helices.^27,61^ Removal of cholesterol from the membrane resulted in an overall reduction of the interactions with SM.

Our analysis indicates a relationship between membrane lipids and specific residues within the transmembrane domain, which are commonly mutated in cancer patients, highlighting their potential involvement in cancer development and altered drug responses (Table S1). Remarkably, some of these mutations change the polarity if even the net charge of the residue. In such cases, the predicted lipid interaction may be particularly disrupted, potentially affecting protein dynamics. It is tempting to speculate how such alteration of protein-lipid interactions could potentially be linked to the onset of the disease.

### Membrane milieu modulates P-gp cavity accessibility

Using MD simulations, we described and compared the structural dynamics of the two conformers, kinked and straight, in a realistic membrane environment. Clear differences between the two conformers were found. Despite both conformers being inward-open, they significantly differ in the volume of the main transmembrane cavity (Fig. 2). P-gp inhibitors bind within this cavity in both conformers, and understanding how this volume changes *in vivo* is crucial for mapping the available “druggable” space, e.g., for developing P-gp inhibitors. The kinked conformer exhibits a much more compact cavity compared to the straight conformer. During the MD simulations, this volume changes, especially with variations in the membrane composition. Specifically, the absence of ATP in the binding site resulted in a compaction of the cavity volume in both conformers (Fig. 2a).

As mentioned above, P-gp also functions as a lipid floppase.^7–9^ Consistent with previous studies,^11,60^ lipids and cholesterol were observed to enter the main cavity during simulations. However, this phenomenon was only observed in the straight conformation (Fig. 2c, d). In contrast, the bending of helices TM4 and TM10 resulted in the blockage of the substrate cavity in the kinked conformation, a characteristic observed across all simulated systems. The accessibility profile of the P-gp substrate cavity is intuitively linked to the aperture angle of its transmembrane helical bundles and, consequently, to the NBD-NBD distance. For instance, in a recent study by Jorgensen and coworkers, a critical aperture angle of 27^◦^ was identified for the entrance of rhodamine.^66^ The conformational dynamics and structural flexibility of the inward-open state of P-gp were further explored using EPR spectroscopy, closely complementing simulations. This highlighted that the distance between the two NBDs could extend beyond the values reported by other structural data, reaching at least 20 Å.^59^

This work demonstrates that the values reported above are frequently observed for both inward-open conformers. However, in our simulations, having a wide angle or a large NBD-NBD distance does not seem to guarantee substrate entrance if helices TM4 and TM10 are kinked.

The unbiased MD simulations sampled the spontaneous transition to an intermediate inward-open state, with TM10 changing from a straight to a kinked conformation. This transition was observed in a particular system (apoSnc) where the membrane lacked cholesterol, potentially resulting in increased membrane fluidity that might have accelerated this transition. Additionally, although the protein was not loaded with nucleotide, it’s plausible that the presence of ATP could influence this transition. Moreover, the order of events, that is, whether TM4 or the (apparently more labile) TM10 kinks first in a functional working cycle, remains an open question (see below).

The observed movement of TM10 could potentially act as a piston mechanism, facilitating the movement of substrates into the transmembrane cavity and subsequent substrate exclusion. Furthermore, the degree of bending may vary to accommodate substrates of different sizes and shapes. Interestingly, membrane components were not observed to enter the cavity from the TM10/TM12 portal, but they could access the passage available from the opposite TM4/TM6 portal. The latter was suggested to be an energetically favorable entrance gate.^63^

It has been suggested that TM4 and TM10 undergo structural rearrangements between straight and kinked conformations during a productive transport cycle.^35^ These two conformers represent distinct functional states. Our simulations suggest that the straight P-gp structure obtained in detergent is a permeable inward-open state that may precede the kinked structure in the sequence of key conformational states that constitute a functional working cycle (see below and Fig. 6).

**Figure 6:**
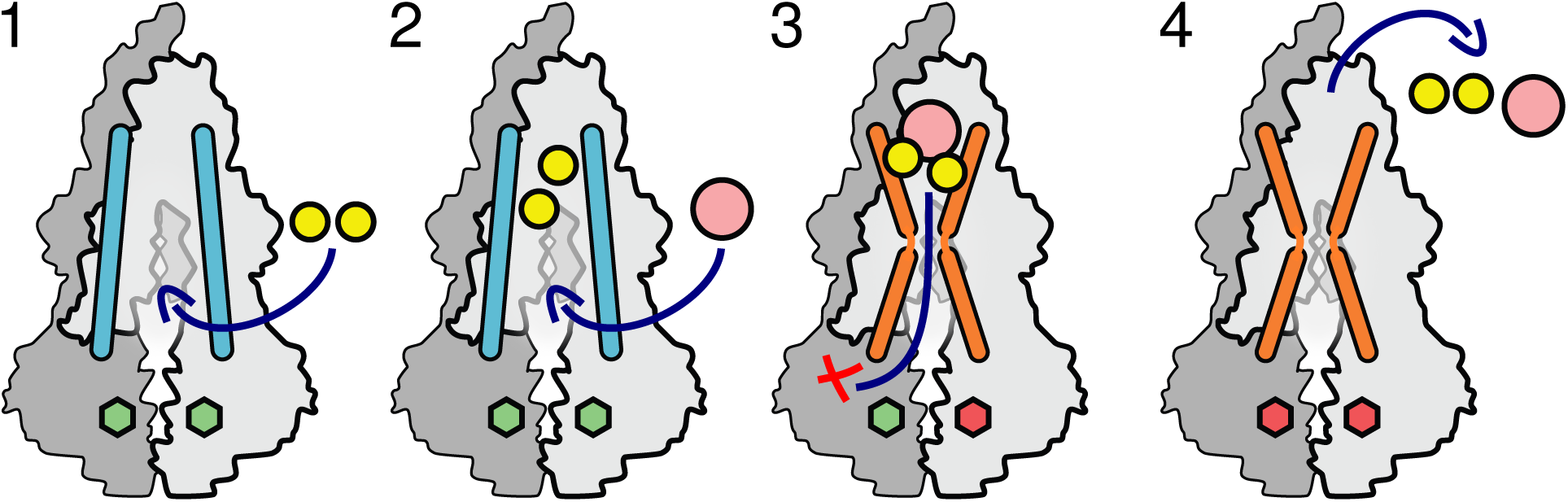
Mechanistic scheme for the straight and kinked inward-open P-gp conformers. P-gp straight (cyan) and kinked (orange) conformers are represented in cartoon. ATP and ADP are shown as green and red hexagons, respectively. Phospholipids and substrate are depicted as yellow and pink spheres, respectively.

### P-gp straight and kinked conformers in a mechanistic context

Lipid snorkeling within the P-gp transmembrane cavity can facilitate the entrance of P-gp substrates. NMR studies have shown that P-gp substrates and modulators bind from the membrane interface region.^10^ MD simulation studies further hypothesized that P-gp inhibitors may promote the recruitment of lipids.^11^ Along the same lines, a pivotal role of lipids in facilitating the binding of the inhibitor tariquidar was hypothesized.^12^ Such co-recruitment has also been discussed for other transporters such as the major facilitator superfamily (MFS) multidrug transporter LmrP from *Lactococcus lactis*, where specific lipids may control the substrate binding pocket, contributing to substrate promiscuity and adaptation.^93^

In light of these evidences, we seek to provide a functional rationale that incorporates the two inward-open conformers. In Fig. 6 a schematic mechanistic timeline is proposed, in the context of a functional cycle. The key steps are as follows. In step 1, P-gp is initially loaded with ATP (Fig. 6, green hexagons) and TM4/TM10 are in the straight conformation. In this resting conformation, lipids (yellow spheres) can enter the transmembrane cavity via both P-gp portals (TM4/TM6 and TM10/TM12), possibly providing a malleable hydrophobic molecular microenvironment, which might facilitate the migration of hydrophobic transport substrates (pink sphere) in step 2. With lipids and substrate present within the main cavity, TM4 and TM10 transition to the kinked conformation (step 3), preventing substrate disengagement. How exactly the protein recognizes whether the substrate is inside the main cavity, and how that signal is transduced to initiate the conformational change of the transmembrane helices, is a formidable question. Possibly, the wrap around the ligand or an engulfed cavity with lipids (or substrate, or both) could shift the conformational equilibrium in favor of a kinked conformation. It is conceivable (but speculative) that one ATP molecule could be hydrolized at this stage to promote the conformational change straight to kinked. In step 4, transport substrate and lipids are finally extruded. ATP hydrolysis then resets the transporter to the initial substrate-receptive state.

## Conclusion

This molecular dynamics (MD) simulation study reinforces and contextualizes the concept that the membrane environment governs the conformational dynamics and flexibility of P-gp. We compare two distinct inward-open conformations, the straight and kinked conformers, with unique structural features and functional implications. The transition between these conformers, particularly the bending and partial unwinding of helices TM4 and TM10, was found to be influenced by membrane composition, including the presence or absence of cholesterol. This work reconciles available structural biology data and sheds light on the potential functional relevance of the two inward-open conformers in the context of the P-gp transport cycle.

## Author Contributions

DD and LVS conceived and designed the research project. DD carried out all simulations, and analyzed and interpreted the data together with LVS. DD and LVS wrote the manuscript.

## Acknowledgment

This work was supported by the German Research Foundation (Deutsche Forschungsgemeinschaft, DFG) through grant SCHA 1574/6-1.

---

**Figure S1.**
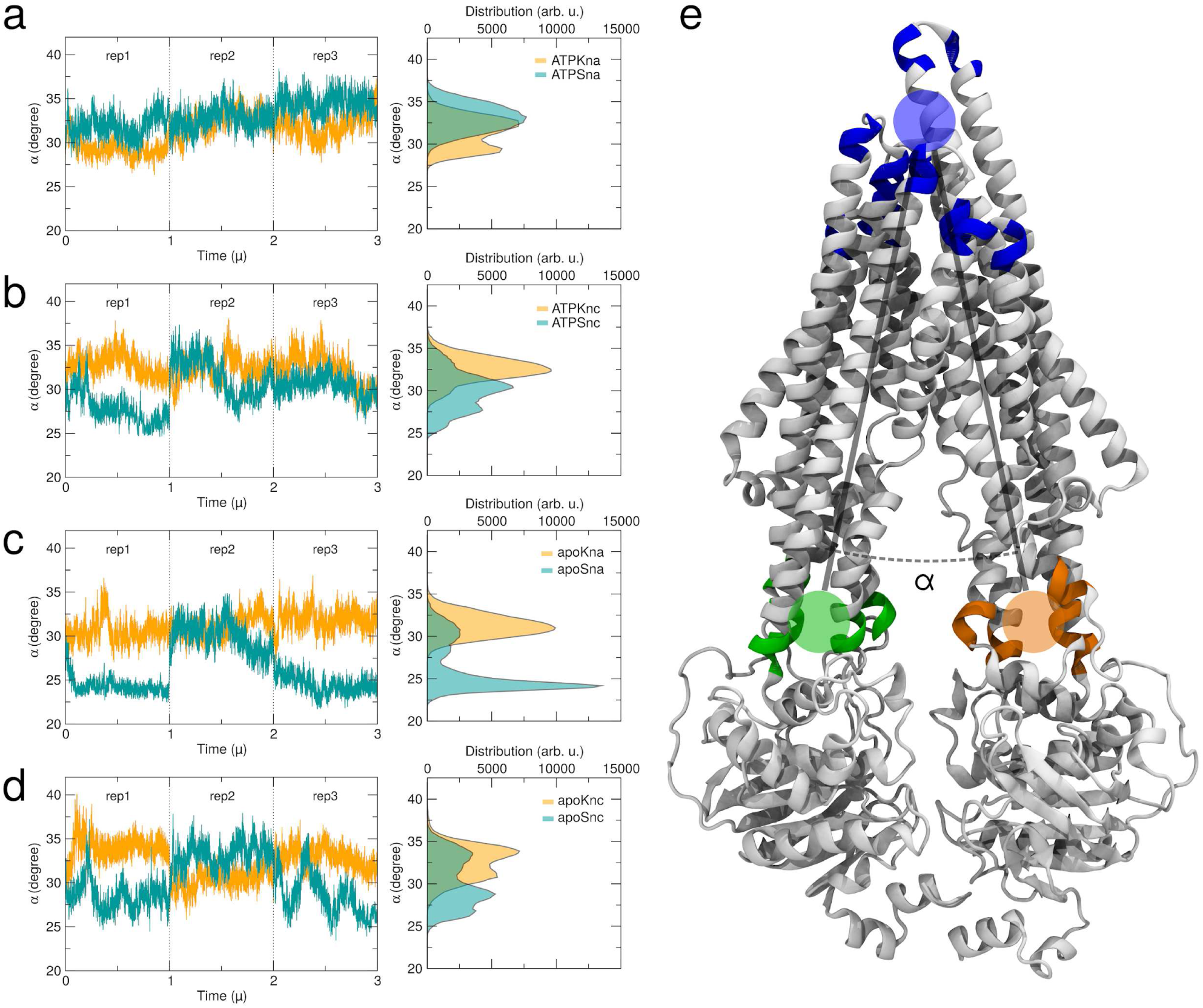
Angle α between the two P-gp transmembrane bundles during simulations. (**a–d**) The time trace of the angle α is shown for each repeat and for each system (the trajectories have been concatenated for clarity). The kinked conformer is indicated in orange and the straight conformer in cyan. The distributions are also shown on the right. (**e**) Ribbon representation of the kinked conformer with the position of the two vectors 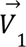 and 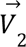 indicated (see Methods).

**Figure S2.**
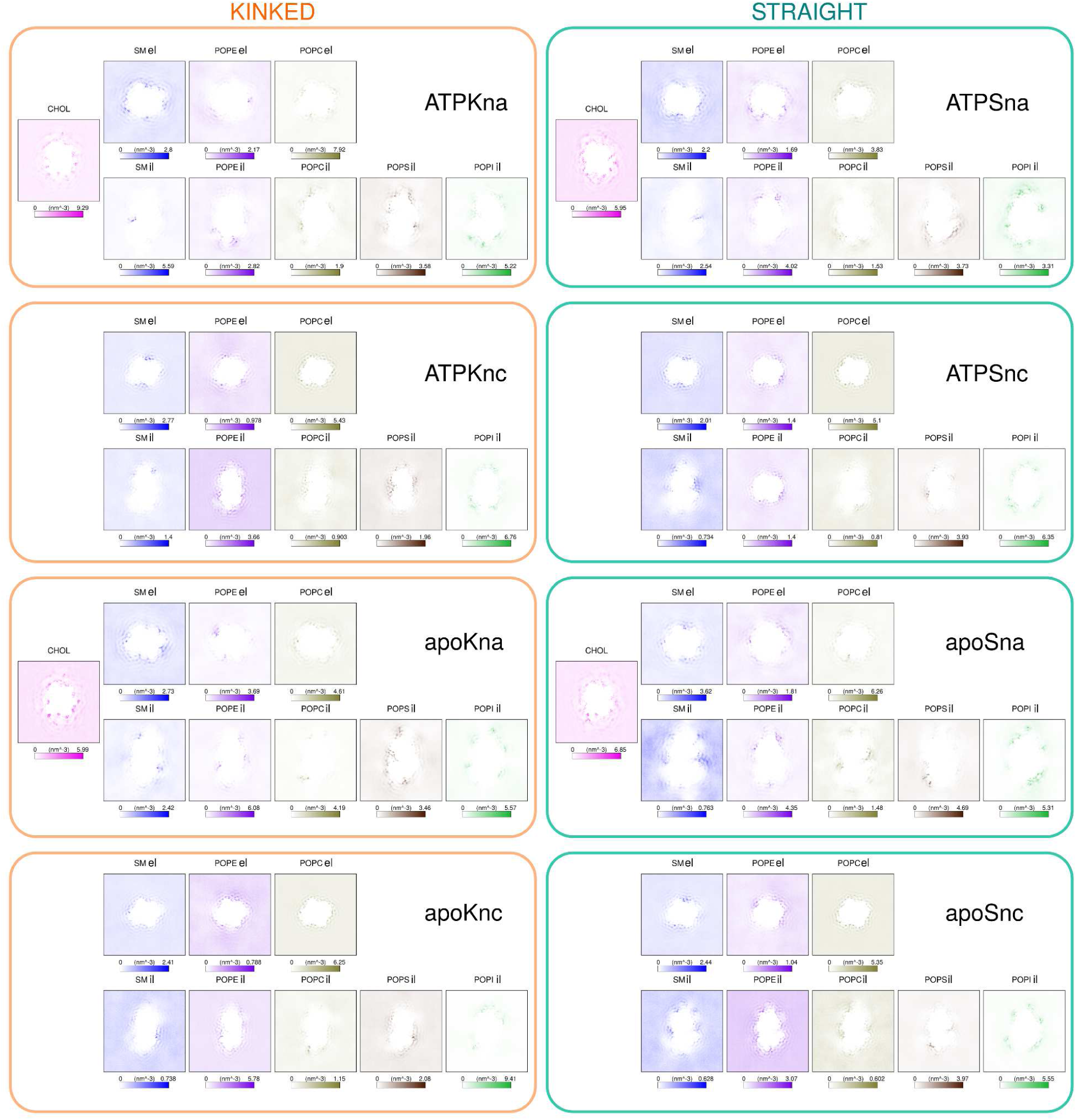
Phospholipids and cholesterol densities from the coarse-grained simulations. The abbreviations for each simulated system are indicated as in Table 1. The color legend for the lipids and cholesterol is the same as in Figure 1. *el*, extracellular leaflet *il*, intracellular leaflet.

**Table S1.**
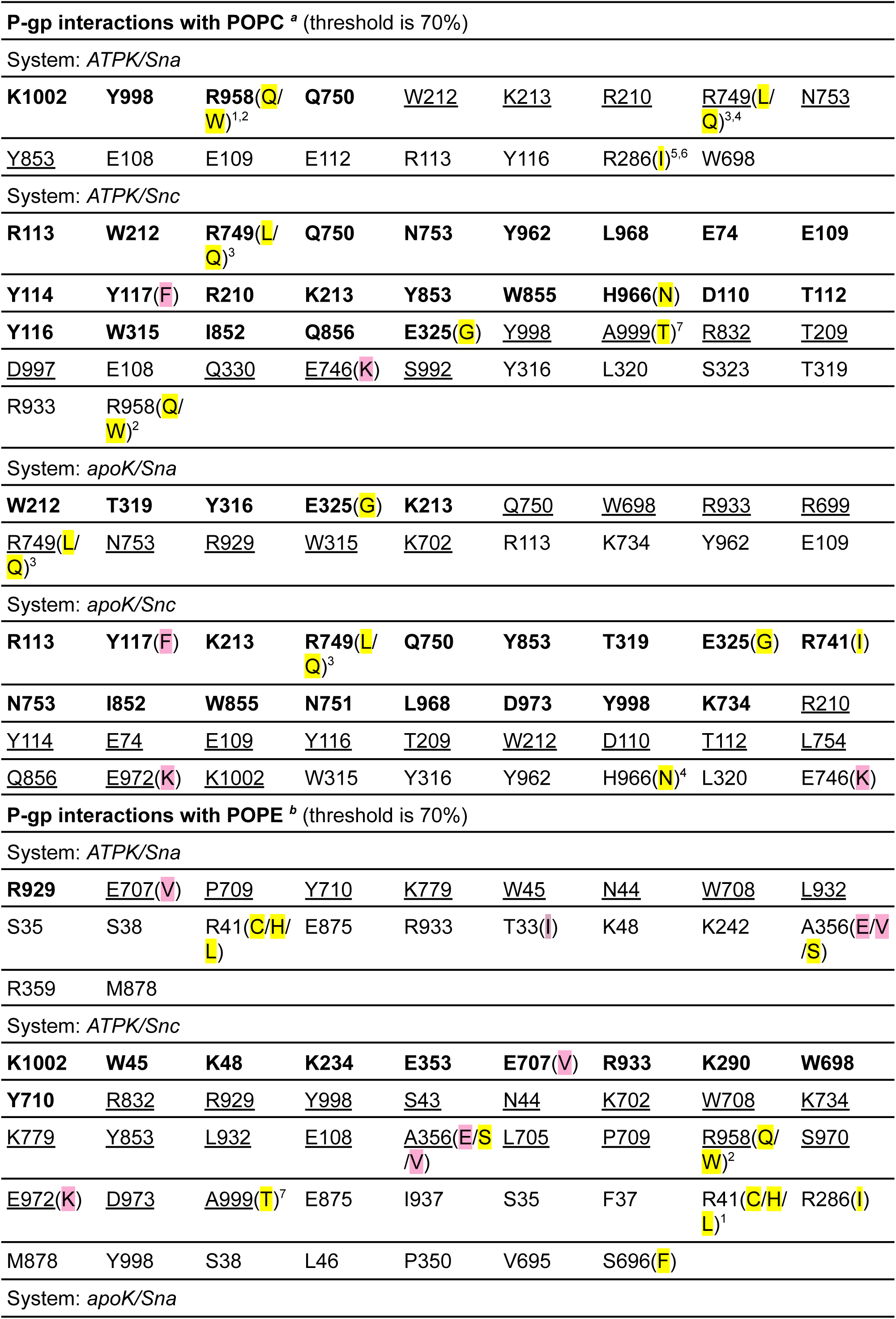
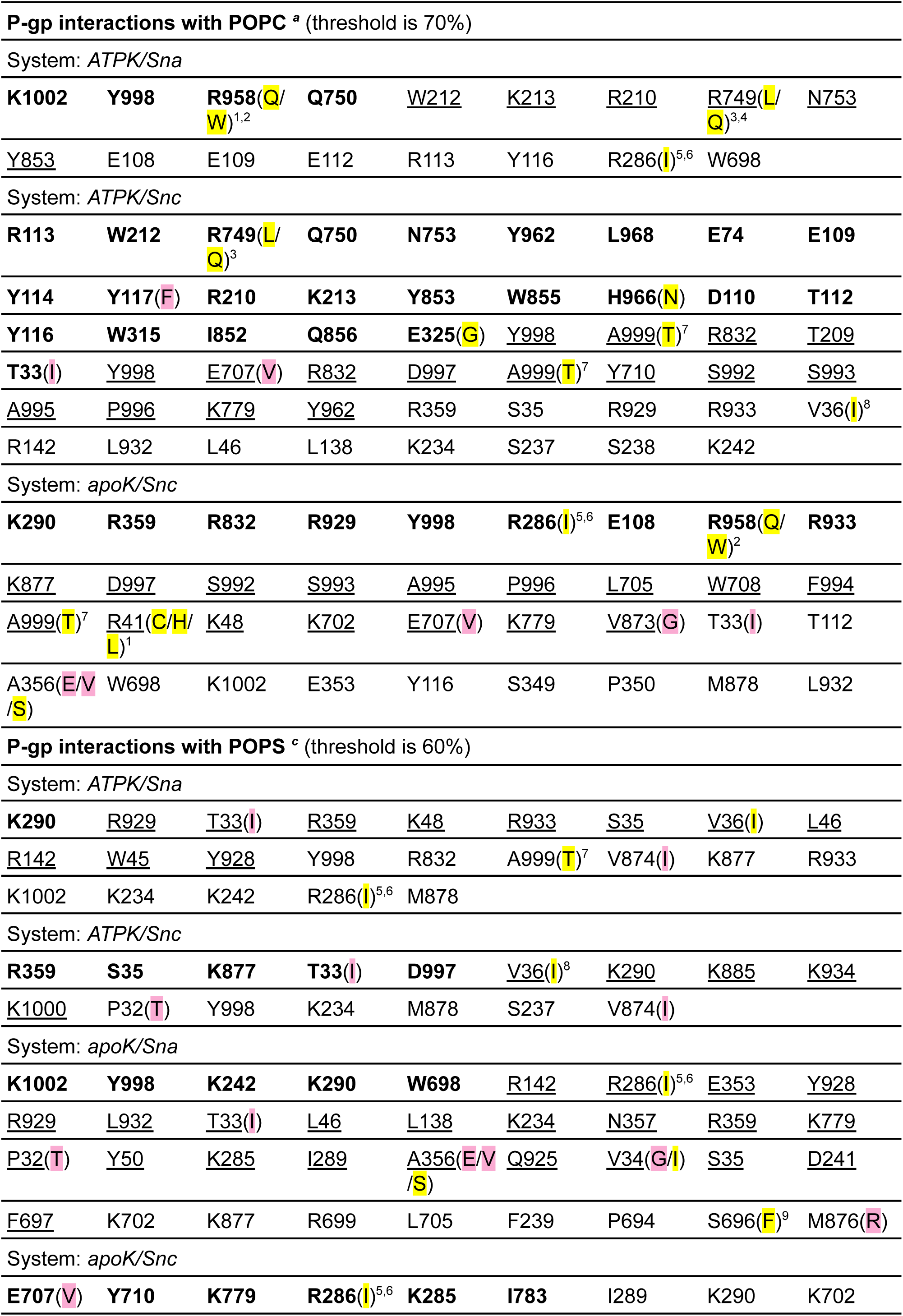
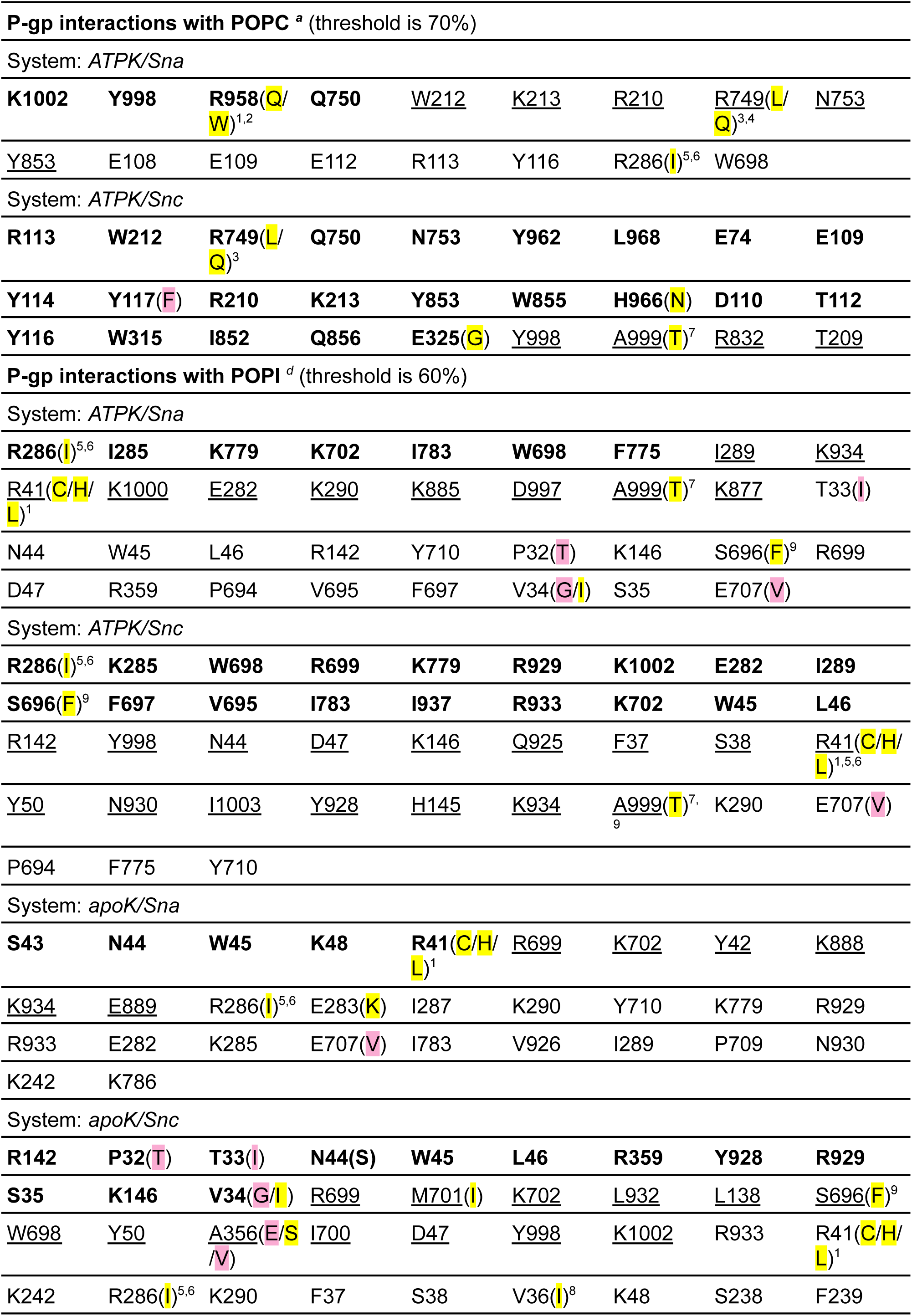
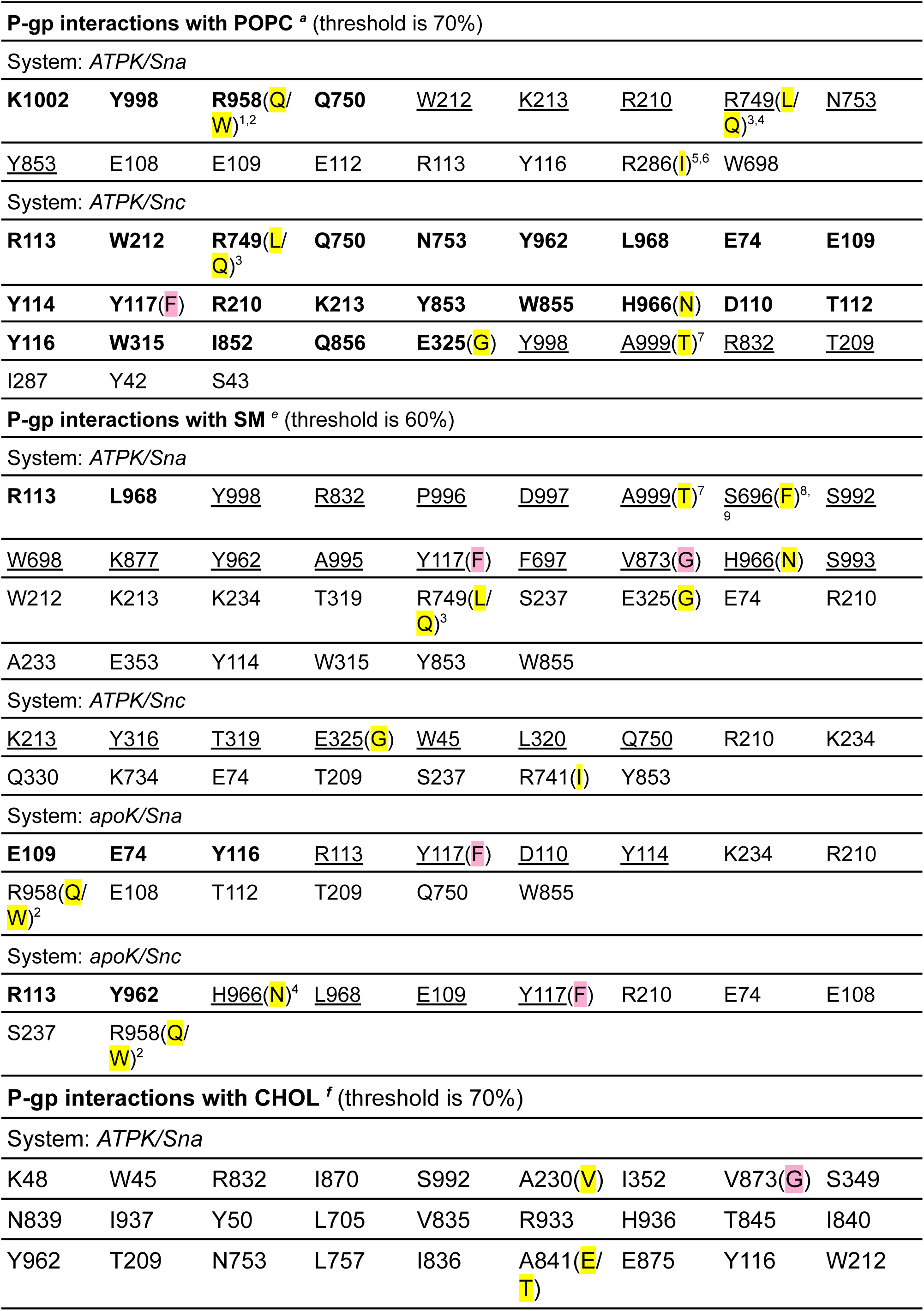
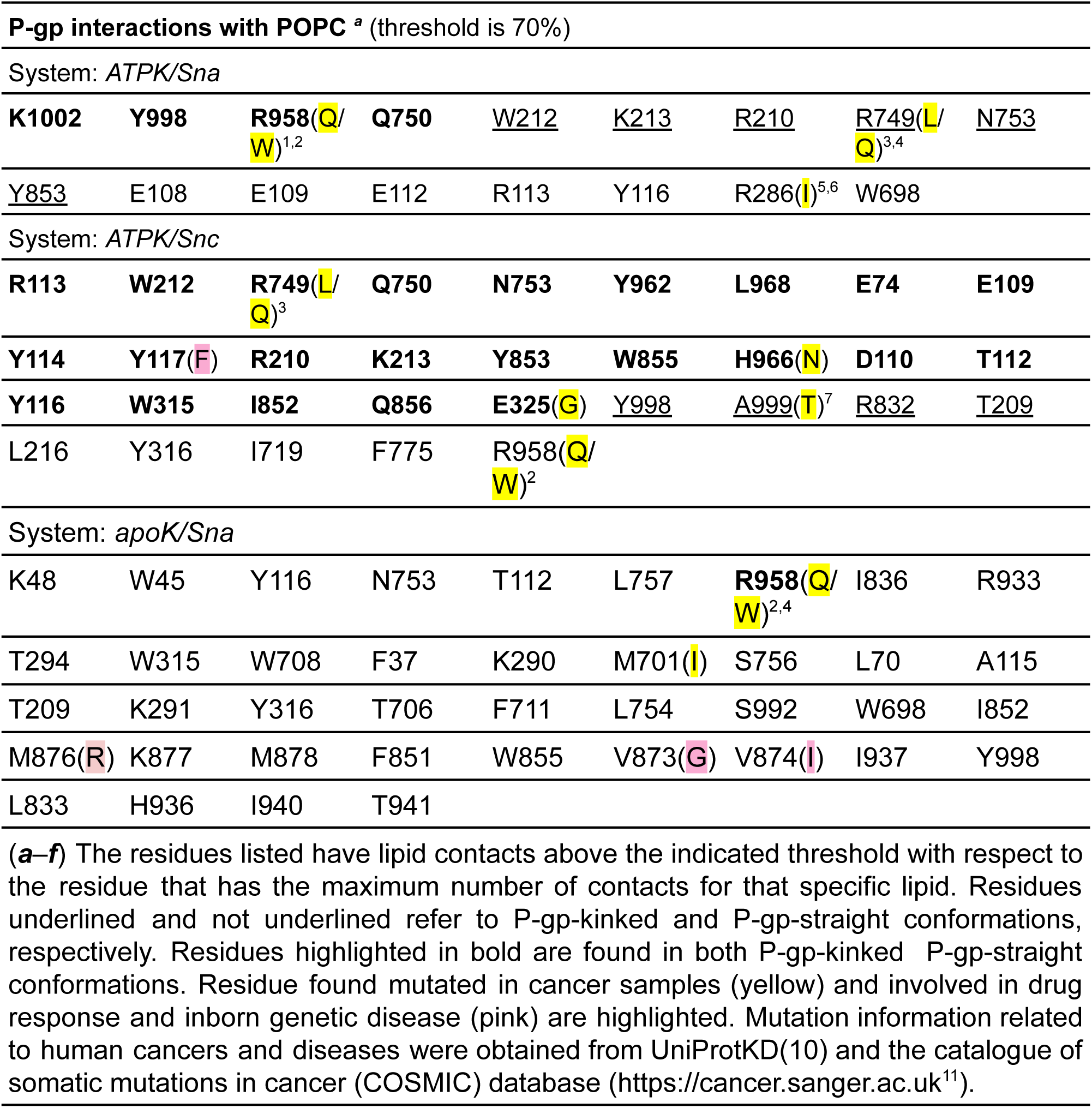
P-gp-lipid and P-gp-cholesterol interactions in all-atom MD simulations.

|  |  |  |  |  |  |  |  |  |
| --- | --- | --- | --- | --- | --- | --- | --- | --- |
| **K1002** | **Y998** | **R958(Q/W)<sup>1,2</sup>** | **Q750** | <u>W212</u> | <u>K213</u> | <u>R210</u> | **R749(L/Q)<sup>3,4</sup>** | <u>N753</u> |
| <u>Y853</u> | <u>E108</u> | <u>E109</u> | <u>E112</u> | <u>R113</u> | <u>Y116</u> | <u>R286(I)<sup>5,6</sup></u> | <u>W698</u> |  |
| System: *ATPK/Snc* |  |  |  |  |  |  |  |  |
| **R113** | **W212** | **R749(L/Q)<sup>3</sup>** | **Q750** | **N753** | **Y962** | **L968** | **E74** | **E109** |
| **Y114** | **Y117(F)** | **R210** | **K213** | **Y853** | **W855** | **H966(N)** | **D110** | **T112** |
| **Y116** | **W315** | **I852** | **Q856** | **E325(G)** | <u>Y998</u> | <u>A999(T)<sup>7</sup></u> | <u>R832</u> | <u>T209</u> |
| <u>D997</u> | <u>E108</u> | <u>Q330</u> | <u>E746(K)</u> | <u>S992</u> | <u>Y316</u> | <u>L320</u> | <u>S323</u> | <u>T319</u> |
| R933 | **R958(Q/W)<sup>2</sup>** |  |  |  |  |  |  |  |
| **W212** | **T319** | **Y316** | **E325(G)** | **K213** | <u>Q750</u> | <u>W698</u> | <u>R933</u> | <u>R699</u> |
| **R749(L/Q)<sup>3</sup>** | <u>N753</u> | <u>R929</u> | <u>W315</u> | <u>K702</u> | <u>R113</u> | <u>K734</u> | <u>Y962</u> | <u>E109</u> |
| System: *apoK/Snc* |  |  |  |  |  |  |  |  |
| **R113** | **Y117(F)** | **K213** | **R749(L/Q)<sup>3</sup>** | **Q750** | **Y853** | **T319** | **E325(G)** | **R741(I)** |
| **N753** | **I852** | **W855** | **N751** | **L968** | **D973** | **Y998** | **K734** | <u>R210</u> |
| <u>Y114</u> | <u>E74</u> | <u>E109</u> | <u>Y116</u> | <u>T209</u> | <u>W212</u> | <u>D110</u> | <u>T112</u> | <u>L754</u> |
| <u>Q856</u> | <u>E972(K)</u> | <u>K1002</u> | <u>W315</u> | <u>Y316</u> | <u>Y962</u> | <u>H966(N)<sup>4</sup></u> | <u>L320</u> | <u>E746(K)</u> |
| **R929** | **E707(V)** | <u>P709</u> | <u>Y710</u> | <u>K779</u> | <u>W45</u> | <u>N44</u> | <u>W708</u> | <u>L932</u> |
| S35 | S38 | **R41(C/H/L)** | <u>E875</u> | <u>R933</u> | <u>T33(I)</u> | <u>K48</u> | <u>K242</u> | **A356(E/V/S)** |
| R359 | M878 |  |  |  |  |  |  |  |
| System: *ATPK/Snc* |  |  |  |  |  |  |  |  |
| **K1002** | **W45** | **K48** | **K234** | **E353** | **E707(V)** | **R933** | **K290** | **W698** |
| **Y710** | <u>R832</u> | <u>R929</u> | <u>Y998</u> | <u>S43</u> | <u>N44</u> | <u>K702</u> | <u>W708</u> | <u>K734</u> |
| <u>K779</u> | <u>Y853</u> | <u>L932</u> | <u>E108</u> | **A356(E/S/V)** | <u>L705</u> | <u>P709</u> | **R958(Q/W)<sup>2</sup>** | <u>S970</u> |
| <u>E972(K)</u> | <u>D973</u> | <u>A999(T)<sup>7</sup></u> | <u>E875</u> | <u>I937</u> | <u>S35</u> | <u>F37</u> | **R41(C/H/L)<sup>1</sup>** | <u>R286(I)</u> |
| M878 | Y998 | S38 | L46 | P350 | V695 | S696(F) |  |  |
| **K1002** | **Y998** | **R958(Q/W)<sup>1,2</sup>** | **Q750** | <u>W212</u> | <u>K213</u> | <u>R210</u> | **R749(L/Q)<sup>3,4</sup>** | <u>N753</u> |
| <u>Y853</u> | <u>E108</u> | <u>E109</u> | <u>E112</u> | <u>R113</u> | <u>Y116</u> | <u>R286(I)<sup>5,6</sup></u> | <u>W698</u> |  |
| System: *ATPK/Snc* |  |  |  |  |  |  |  |  |
| **R113** | **W212** | **R749(L/Q)<sup>3</sup>** | **Q750** | **N753** | **Y962** | **L968** | **E74** | **E109** |
| **Y114** | **Y117(F)** | **R210** | **K213** | **Y853** | **W855** | **H966(N)** | **D110** | **T112** |
| **Y116** | **W315** | **I852** | **Q856** | **E325(G)** | <u>Y998</u> | <u>A999(T)<sup>7</sup></u> | <u>R832</u> | <u>T209</u> |
| **T33(I)** | <u>Y998</u> | <u>E707(V)</u> | <u>R832</u> | <u>D997</u> | <u>A999(T)<sup>7</sup></u> | <u>Y710</u> | <u>S992</u> | <u>S993</u> |
| <u>A995</u> | <u>P996</u> | <u>K779</u> | <u>Y962</u> | <u>R359</u> | <u>S35</u> | <u>R929</u> | <u>R933</u> | <u>V36(I)<sup>8</sup></u> |
| <u>R142</u> | <u>L932</u> | <u>L46</u> | <u>L138</u> | <u>K234</u> | <u>S237</u> | <u>S238</u> | <u>K242</u> |  |
| System: *apoK/Snc* |  |  |  |  |  |  |  |  |
| **K290** | **R359** | **R832** | **R929** | **Y998** | **R286(I)<sup>5,6</sup>** | **E108** | **R958(Q/W)<sup>2</sup>** | **R933** |
| <u>K877</u> | <u>D997</u> | <u>S992</u> | <u>S993</u> | <u>A995</u> | <u>P996</u> | <u>L705</u> | <u>W708</u> | <u>F994</u> |
| <u>A999(T)<sup>7</sup></u> | **R41(C/H/L)<sup>1</sup>** | <u>K48</u> | <u>K702</u> | <u>E707(V)</u> | <u>K779</u> | <u>V873(G)</u> | <u>T33(I)</u> | <u>T112</u> |
| <u>A356(E/V/S)</u> | <u>W698</u> | <u>K1002</u> | <u>E353</u> | <u>Y116</u> | <u>S349</u> | <u>P350</u> | <u>M878</u> | <u>L932</u> |
| **K290** | <u>R929</u> | <u>T33(I)</u> | <u>R359</u> | <u>K48</u> | <u>R933</u> | <u>S35</u> | <u>V36(I)</u> | <u>L46</u> |
| <u>R142</u> | <u>W45</u> | <u>Y928</u> | <u>Y998</u> | <u>R832</u> | <u>A999(T)<sup>7</sup></u> | <u>V874(I)</u> | <u>K877</u> | <u>R933</u> |
| <u>K1002</u> | <u>K234</u> | <u>K242</u> | <u>R286(I)<sup>5,6</sup></u> | <u>M878</u> |  |  |  |  |
| System: *ATPK/Snc* |  |  |  |  |  |  |  |  |
| **R359** | **S35** | **K877** | **T33(I)** | **D997** | <u>V36(I)<sup>8</sup></u> | <u>K290</u> | <u>K885</u> | <u>K934</u> |
| <u>K1000</u> | <u>P32(T)</u> | <u>Y998</u> | <u>K234</u> | <u>M878</u> | <u>S237</u> | <u>V874(I)</u> |  |  |
| **K1002** | **Y998** | **K242** | **K290** | **W698** | <u>R142</u> | <u>R286(I)<sup>5,6</sup></u> | <u>E353</u> | <u>Y928</u> |
| <u>R929</u> | <u>L932</u> | <u>T33(I)</u> | <u>L46</u> | <u>L138</u> | <u>K234</u> | <u>N357</u> | <u>R359</u> | <u>K779</u> |
| <u>P32(T)</u> | <u>Y50</u> | <u>K285</u> | <u>I289</u> | <u>A356(E/V/S)</u> | <u>Q925</u> | <u>V34(G/I)</u> | <u>S35</u> | <u>D241</u> |
| <u>F697</u> | <u>K702</u> | <u>K877</u> | <u>R699</u> | <u>L705</u> | <u>F239</u> | <u>P694</u> | <u>S696(F)<sup>9</sup></u> | <u>M876(R)</u> |
| System: *apoK/Snc* |  |  |  |  |  |  |  |  |
| **E707(V)** | **Y710** | **K779** | **R286(I)<sup>5,6</sup>** | **K285** | **I783** | <u>I289</u> | <u>K290</u> | <u>K702</u> |
| **K1002** | **Y998** | **R958**(Q/<br>W)<sup>1,2</sup> | **Q750** | <u>W212</u> | <u>K213</u> | <u>R210</u> | <u>R749</u>(L/<br>Q)<sup>3,4</sup> | <u>N753</u> |
| <u>Y853</u> | <u>E108</u> | <u>E109</u> | <u>E112</u> | <u>R113</u> | <u>Y116</u> | <u>R286</u>(I)<sup>5,6</sup> | <u>W698</u> |  |
| System: *ATPK/Snc* |  |  |  |  |  |  |  |  |
| **R113** | **W212** | **R749**(L/<br>Q)<sup>3</sup> | **Q750** | **N753** | **Y962** | **L968** | **E74** | **E109** |
| **Y114** | **Y117**(F) | **R210** | **K213** | **Y853** | **W855** | **H966**(N) | **D110** | **T112** |
| **Y116** | **W315** | **I852** | **Q856** | **E325**(G) | <u>Y998</u> | <u>A999</u>(T)<sup>7</sup> | <u>R832</u> | <u>T209</u> |
| **R286**(I)<sup>5,6</sup> | **I285** | **K779** | **K702** | **I783** | **W698** | **F775** | <u>I289</u> | <u>K934</u> |
| <u>R41</u>(C/H/<br>L)<sup>1</sup> | <u>K1000</u> | <u>E282</u> | <u>K290</u> | <u>K885</u> | <u>D997</u> | <u>A999</u>(T)<sup>7</sup> | <u>K877</u> | <u>T33</u>(I) |
| <u>N44</u> | <u>W45</u> | <u>L46</u> | <u>R142</u> | <u>Y710</u> | <u>P32</u>(T) | <u>K146</u> | <u>S696</u>(F)<sup>9</sup> | <u>R699</u> |
| <u>D47</u> | <u>R359</u> | <u>P694</u> | <u>V695</u> | <u>F697</u> | <u>V34</u>(G/I) | <u>S35</u> | <u>E707</u>(V) |  |
| System: *ATPK/Snc* |  |  |  |  |  |  |  |  |
| **R286**(I)<sup>5,6</sup> | **K285** | **W698** | **R699** | **K779** | **R929** | **K1002** | **E282** | **I289** |
| **S696**(F)<sup>9</sup> | **F697** | **V695** | **I783** | **I937** | **R933** | **K702** | **W45** | **L46** |
| <u>R142</u> | <u>Y998</u> | <u>N44</u> | <u>D47</u> | <u>K146</u> | <u>Q925</u> | <u>F37</u> | <u>S38</u> | <u>R41</u>(C/H/<br>L)<sup>1,5,6</sup> |
| <u>Y50</u> | <u>N930</u> | <u>I1003</u> | <u>Y928</u> | <u>H145</u> | <u>K934</u> | <u>A999</u>(T)<sup>7,9</sup> | <u>K290</u> | <u>E707</u>(V) |
| <u>P694</u> | <u>F775</u> | <u>Y710</u> |  |  |  |  |  |  |
| **S43** | **N44** | **W45** | **K48** | **R41**(C/H/<br>L)<sup>1</sup> | <u>R699</u> | <u>K702</u> | <u>Y42</u> | <u>K888</u> |
| <u>K934</u> | <u>E889</u> | <u>R286</u>(I)<sup>5,6</sup> | <u>E283</u>(K) | <u>I287</u> | <u>K290</u> | <u>Y710</u> | <u>K779</u> | <u>R929</u> |
| <u>R933</u> | <u>E282</u> | <u>K285</u> | <u>E707</u>(V) | <u>I783</u> | <u>V926</u> | <u>I289</u> | <u>P709</u> | <u>N930</u> |
| <u>K242</u> | <u>K786</u> |  |  |  |  |  |  |  |
| System: *apoK/Snc* |  |  |  |  |  |  |  |  |
| **R142** | **P32**(T) | **T33**(I) | **N44**(S) | **W45** | **L46** | **R359** | **Y928** | **R929** |
| **S35** | **K146** | **V34**(G/I) | <u>R699</u> | <u>M701</u>(I) | <u>K702</u> | <u>L932</u> | <u>L138</u> | <u>S696</u>(F)<sup>9</sup> |
| <u>W698</u> | <u>Y50</u> | <u>A356</u>(E/S<br>/V) | <u>I700</u> | <u>D47</u> | <u>Y998</u> | <u>K1002</u> | <u>R933</u> | <u>R41</u>(C/H/<br>L)<sup>1</sup> |
| <u>K242</u> | <u>R286</u>(I)<sup>5,6</sup> | <u>K290</u> | <u>F37</u> | <u>S38</u> | <u>V36</u>(I)<sup>8</sup> | <u>K48</u> | <u>S238</u> | <u>F239</u> |
| **K1002** | **Y998** | **R958(Q/W)<sup>1,2</sup>** | **Q750** | <u>W212</u> | <u>K213</u> | <u>R210</u> | **R749(L/Q)<sup>3,4</sup>** | <u>N753</u> |
| <u>Y853</u> | E108 | E109 | E112 | R113 | Y116 | R286(I)<sup>5,6</sup> | W698 |  |
| System: *ATPK/Snc* |  |  |  |  |  |  |  |  |
| **R113** | **W212** | **R749(L/Q)<sup>3</sup>** | **Q750** | **N753** | **Y962** | **L968** | **E74** | **E109** |
| **Y114** | **Y117(F)** | **R210** | **K213** | **Y853** | **W855** | **H966(N)** | **D110** | **T112** |
| **Y116** | **W315** | **I852** | **Q856** | **E325(G)** | <u>Y998</u> | <u>A999(T)<sup>7</sup></u> | <u>R832</u> | <u>T209</u> |
| I287 | Y42 | S43 |  |  |  |  |  |  |
| **R113** | **L968** | <u>Y998</u> | <u>R832</u> | <u>P996</u> | <u>D997</u> | <u>A999(T)<sup>7</sup></u> | <u>S696(F)<sup>8,9</sup></u> | <u>S992</u> |
| <u>W698</u> | <u>K877</u> | <u>Y962</u> | <u>A995</u> | <u>Y117(F)</u> | <u>F697</u> | <u>V873(G)</u> | <u>H966(N)</u> | <u>S993</u> |
| W212 | K213 | K234 | T319 | **R749(L/Q)<sup>3</sup>** | S237 | E325(G) | E74 | R210 |
| A233 | E353 | Y114 | W315 | Y853 | W855 |  |  |  |
| System: *ATPK/Snc* |  |  |  |  |  |  |  |  |
| <u>K213</u> | <u>Y316</u> | <u>T319</u> | <u>E325(G)</u> | <u>W45</u> | <u>L320</u> | <u>Q750</u> | R210 | K234 |
| Q330 | K734 | E74 | T209 | S237 | R741(I) | Y853 |  |  |
| **E109** | **E74** | **Y116** | <u>R113</u> | <u>Y117(F)</u> | <u>D110</u> | <u>Y114</u> | K234 | R210 |
| **R958(Q/W)<sup>2</sup>** | E108 | T112 | T209 | Q750 | W855 |  |  |  |
| System: *apoK/Snc* |  |  |  |  |  |  |  |  |
| **R113** | **Y962** | <u>H966(N)<sup>4</sup></u> | <u>L968</u> | <u>E109</u> | <u>Y117(F)</u> | R210 | E74 | E108 |
| S237 | **R958(Q/W)<sup>2</sup>** |  |  |  |  |  |  |  |
| K48 | W45 | R832 | I870 | S992 | **A230(V)** | I352 | **V873(G)** | S349 |
| N839 | I937 | Y50 | L705 | V835 | R933 | H936 | T845 | I840 |
| Y962 | T209 | N753 | L757 | I836 | **A841(E/T)** | E875 | Y116 | W212 |

**Table S1. P-gp-lipid and P-gp-cholesterol interactions in all-atom MD simulations.****P-gp interactions with POPC <sup>a</sup> (threshold is 70%)**System: *ATPK/Sna*
|  |  |  |  |  |  |  |  |  |
| --- | --- | --- | --- | --- | --- | --- | --- | --- |
| **K1002** | **Y998** | **R958(Q/W)<sup>1,2</sup>** | **Q750** | <u>W212</u> | <u>K213</u> | <u>R210</u> | <u>R749(L/Q)<sup>3,4</sup></u> | <u>N753</u> |
| <u>Y853</u> | E108 | E109 | E112 | R113 | Y116 | R286(I)<sup>5,6</sup> | W698 |  |
| System: *ATPK/Snc* |  |  |  |  |  |  |  |  |
| **R113** | **W212** | **R749(L/Q)<sup>3</sup>** | **Q750** | **N753** | **Y962** | **L968** | **E74** | **E109** |
| **Y114** | **Y117(F)** | **R210** | **K213** | **Y853** | **W855** | **H966(N)** | **D110** | **T112** |
| **Y116** | **W315** | **I852** | **Q856** | **E325(G)** | <u>Y998</u> | <u>A999(T)<sup>7</sup></u> | <u>R832</u> | <u>T209</u> |
| L216 | Y316 | I719 | F775 | R958(Q/W)<sup>2</sup> |  |  |  |  |
| K48 | W45 | Y116 | N753 | T112 | L757 | **R958(Q/W)<sup>2,4</sup>** | I836 | R933 |
| T294 | W315 | W708 | F37 | K290 | M701(I) | S756 | L70 | A115 |
| T209 | K291 | Y316 | T706 | F711 | L754 | S992 | W698 | I852 |
| M876(R) | K877 | M878 | F851 | W855 | V873(G) | V874(I) | I937 | Y998 |
| L833 | H936 | I940 | T941 |  |  |  |  |  |

(a–f) The residues listed have lipid contacts above the indicated threshold with respect to the residue that has the maximum number of contacts for that specific lipid. Residues underlined and not underlined refer to P-gp-kinked and P-gp-straight conformations, respectively. Residues highlighted in bold are found in both P-gp-kinked P-gp-straight conformations. Residue found mutated in cancer samples (yellow) and involved in drug response and inborn genetic disease (pink) are highlighted. Mutation information related to human cancers and diseases were obtained from UniProtKD(10) and the catalogue of somatic mutations in cancer (COSMIC) database (https://cancer.sanger.ac.uk<sup>11</sup>).

**Figure S3.**
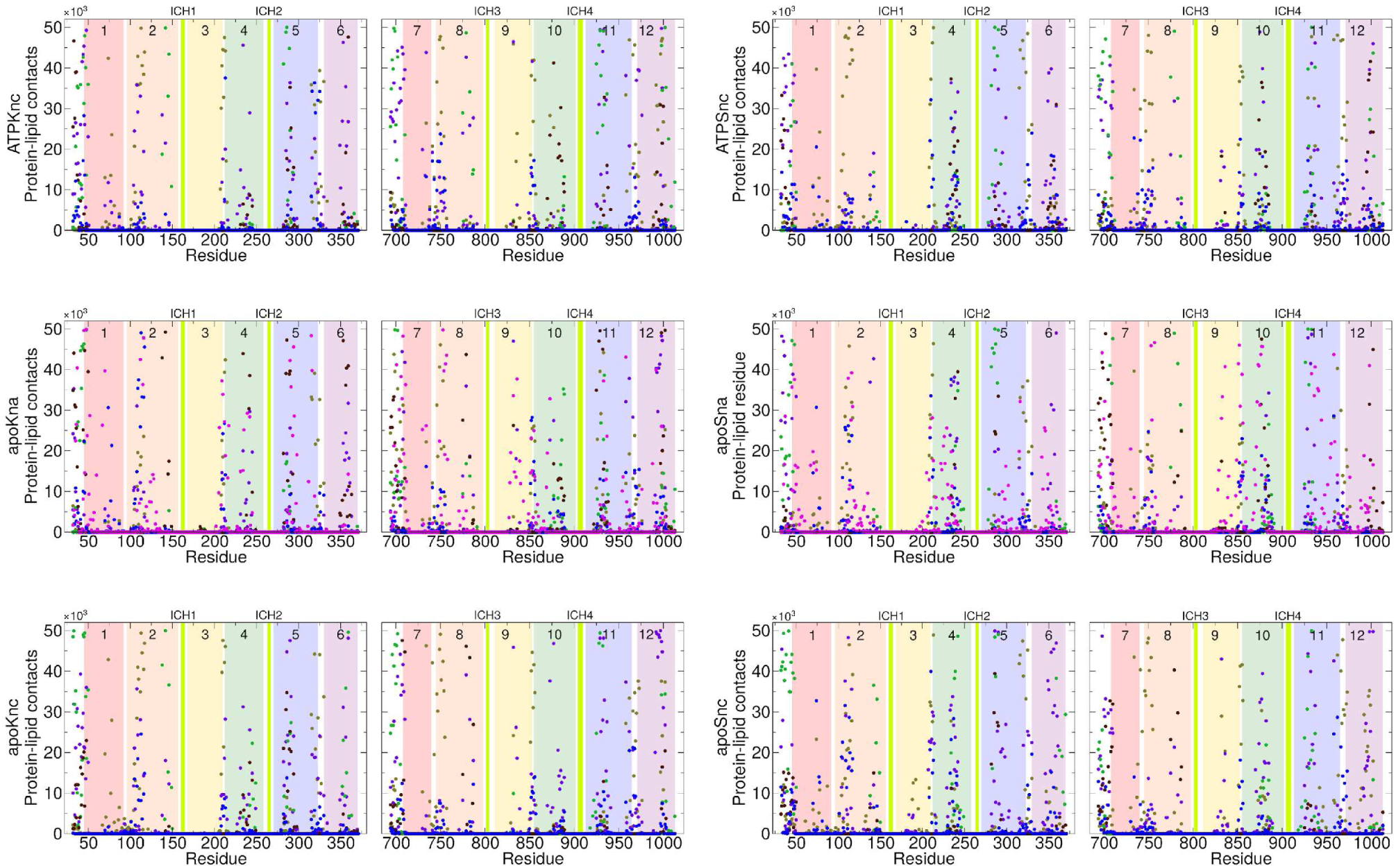
Protein-lipid contact analysis of the simulated P-gp conformers. Overall P-gp/lipid and P-gp/cholesterol contacts in the 3 μs of collected simulation time of all the simulated systems for TM1 to TM6 (residues 22–381) and TM7 to TM12 (residues 684–1023). The P-gp transmembrane helices are numbered and colored in rainbow as in Fig. 1. The color legend for the lipids and cholesterol is the same as in Fig. 1 in the main text. Intracellular coupling helices ICH1 to ICH4 are indicated in green.

**Figure S4.**
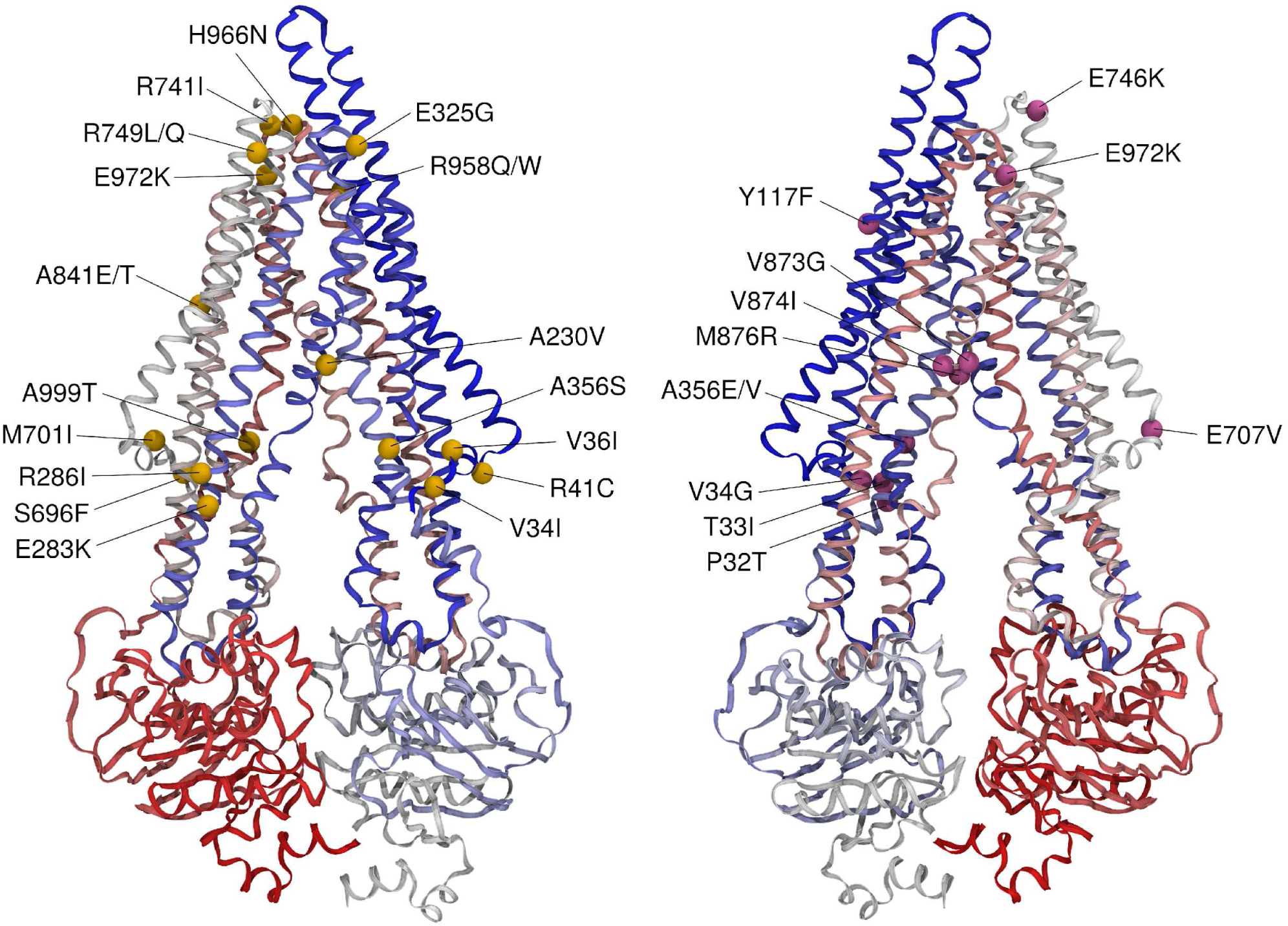
P-gp residues mutated in cancer and found in contact with membrane lipids. The Cα of mutated residues are indicated. The color code is the same as in Table S1. In yellow (left) residues found in cancer cells, in pink (right) residues involved in drug response (tramadol response) and inborn genetic disease. Mutation information related to human cancers and diseases were obtained from the UniProtKB(10) and the catalogue of somatic mutations in cancer (COSMIC) database (https://cancer.sanger.ac.uk)(11).

**Figure S5.**
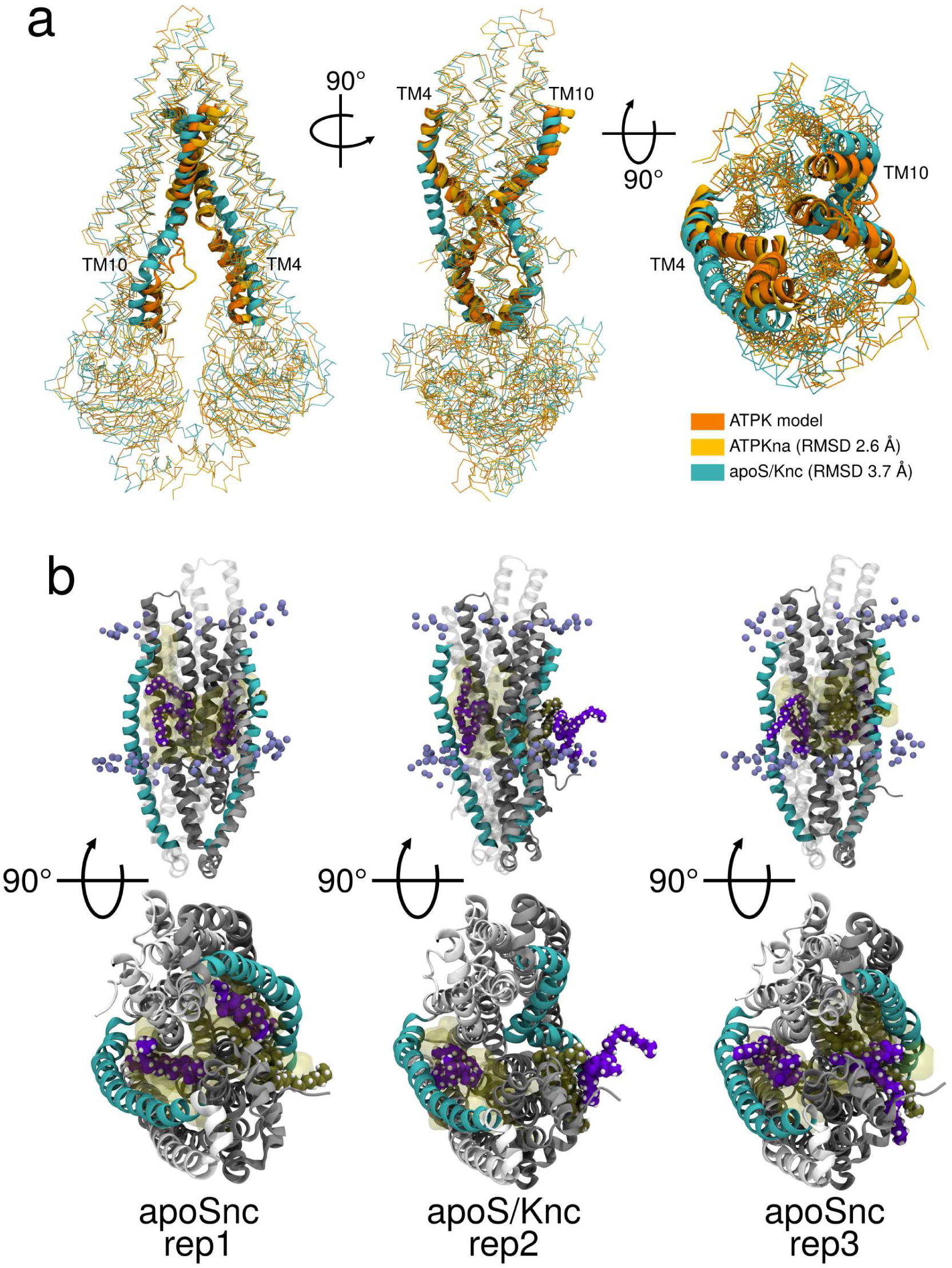
The half-straight/half-kinked P-gp conformer. Structural comparison between the initial P-gp kinked conformer (orange), the simulated P-gp kinked (light orange) and the simulated straight (cyan) conformers. The simulated P-gp kinked conformer shown (light orange) is the last frame from the “repeat 1” simulation of the ATPKna system. The P-gp straight conformer shown (cyan) is a frame from the “repeat 2” simulation of the apoSnc system. (**a**) The TM4 and TM10 are shown in ribbon, the rest of the protein as Cα trace. Structures have been superposed over the Cα atoms of the transmembrane domain (residues 32–371 and 694–1013), the RMSD is indicated. (**b**) Snapshots from the apoSnc simulated systems. The repeat is indicated, showing the transmembrane cavity of the P-gp straight conformer with the membrane components (in van der Waals representation), snorkeling within the cavity (indicated in yellow). The phosphorus atoms from the lipid headgroups are indicated as spheres. Solvent, ions and NBD domains are not shown for clarity.

**Figure S6.**
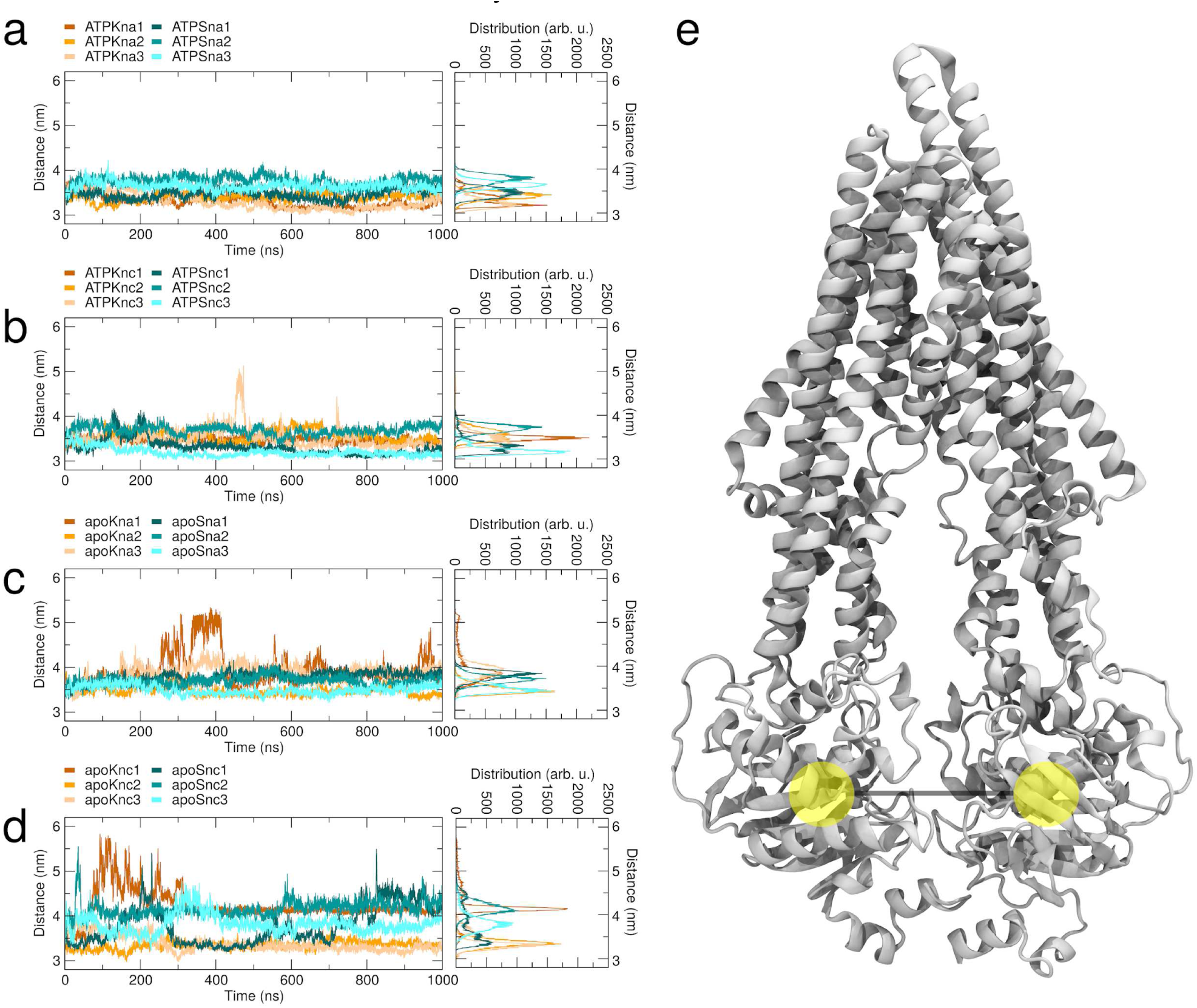
Distance between the center of mass of the two P-gp NBDs during simulations. (**a–d**) The time trace of the distance is shown for each repeat simulation (1,2,3) and for each conformer. The kinked conformer is indicated in orange and the straight conformer in cyan. The distributions are also shown on the right. (**e**) Ribbon representation of the kinked conformer with the position of the NBD centre of masses highlighted in yellow.

**Figure S7.**
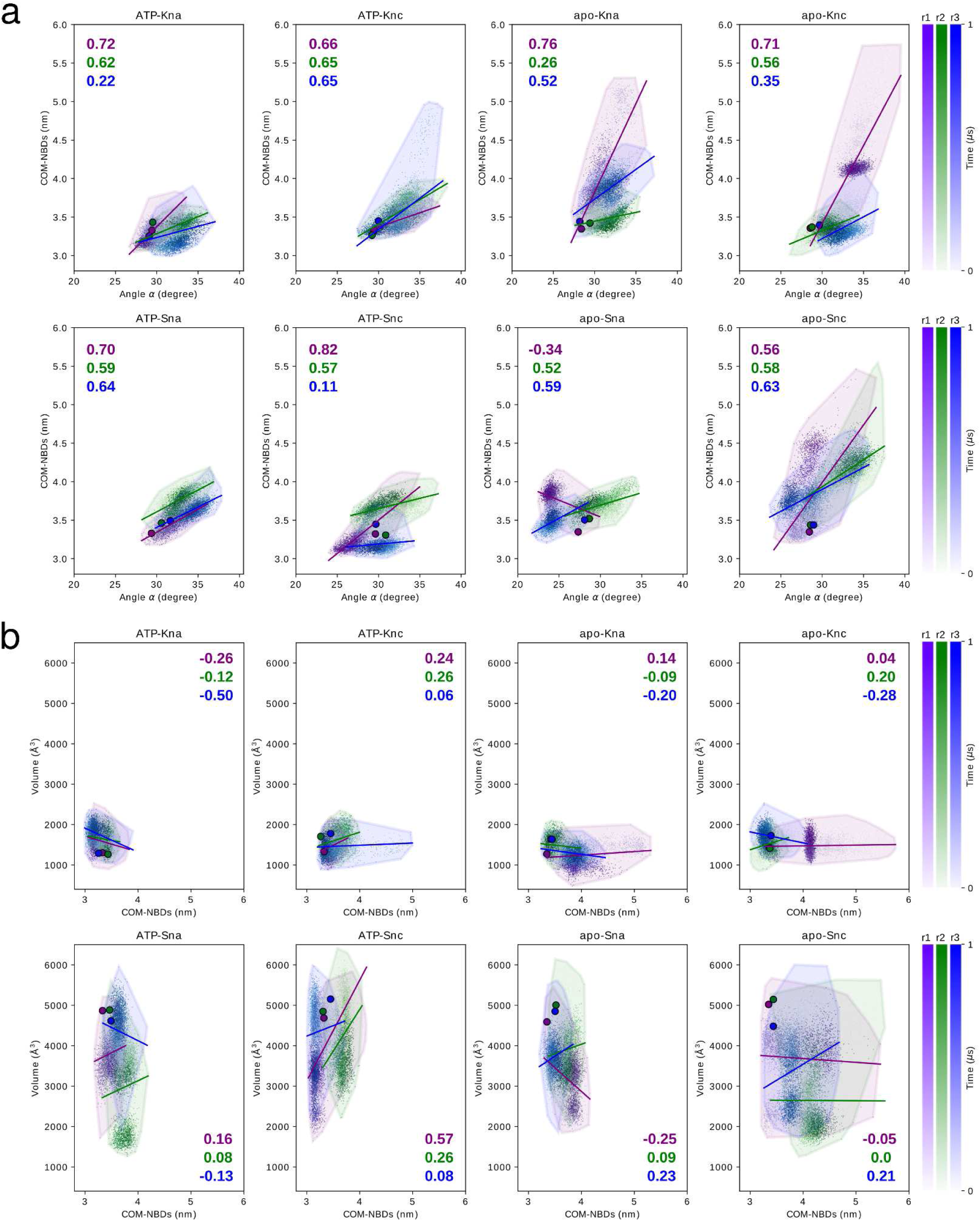
Structural correlation of P-gp. (**a**) Correlation between the NBD dimer distance and the angle α between the transmembrane bundles. (**b**) Correlation between the volume and the NBD dimer distance. For each simulated system and each repeat, the Pearson correlation coefficients are reported. The three simulation repeats are indicated in different colors, with the color gradient indicating the simulation time as in Fig. 4 in the main text. The circles are the initial values at the beginning of the simulations.

**Figure S8.**
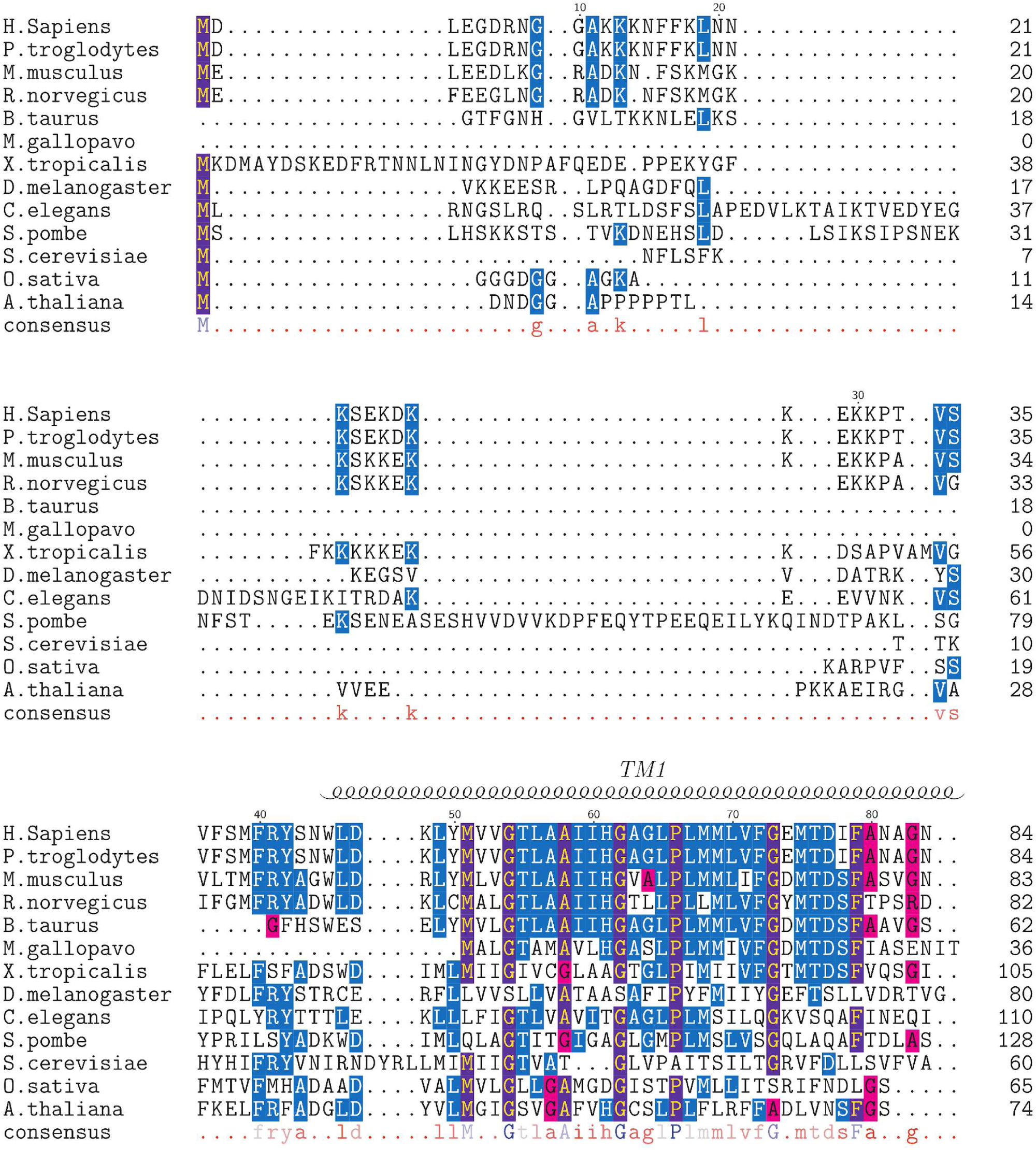
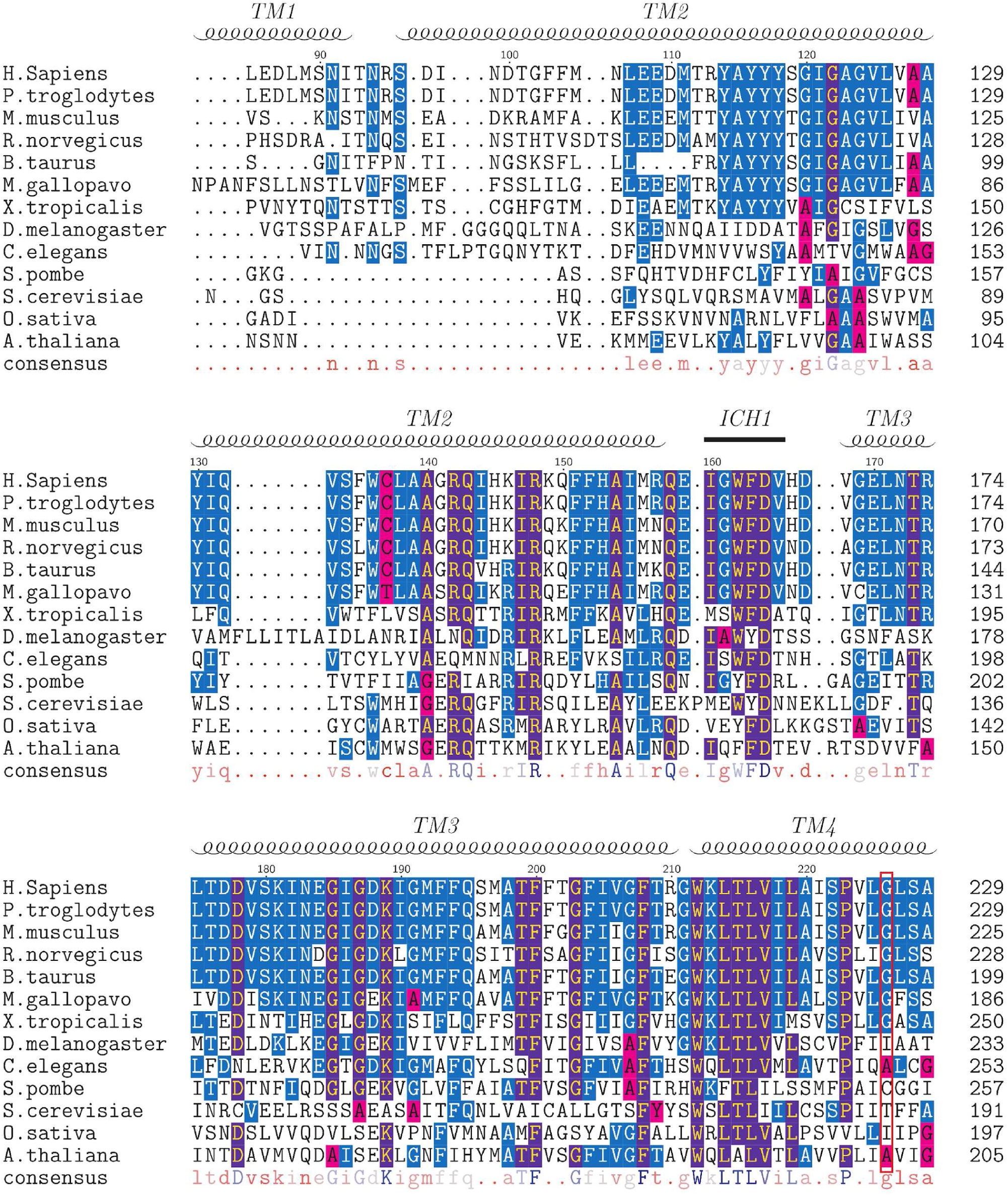
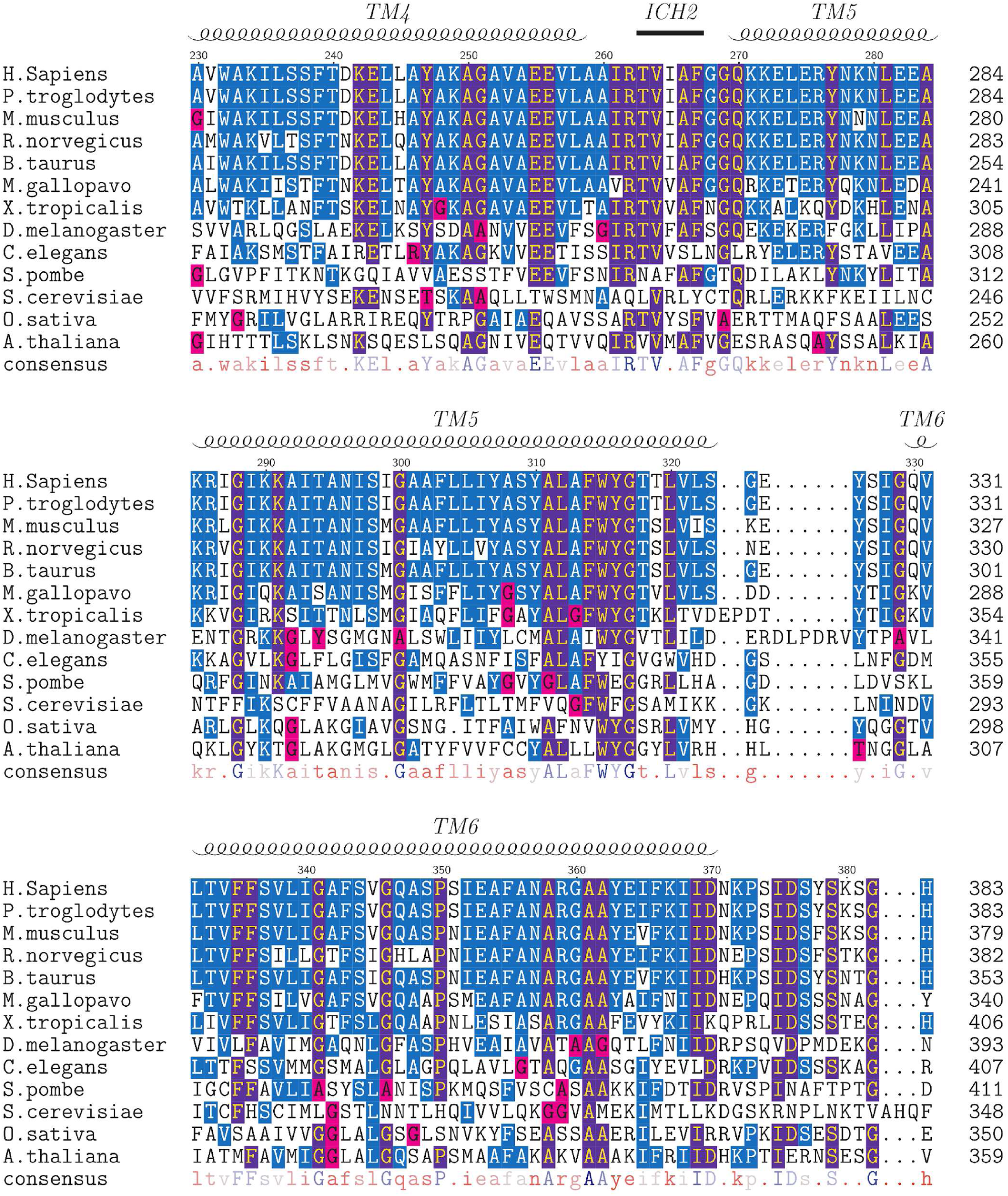
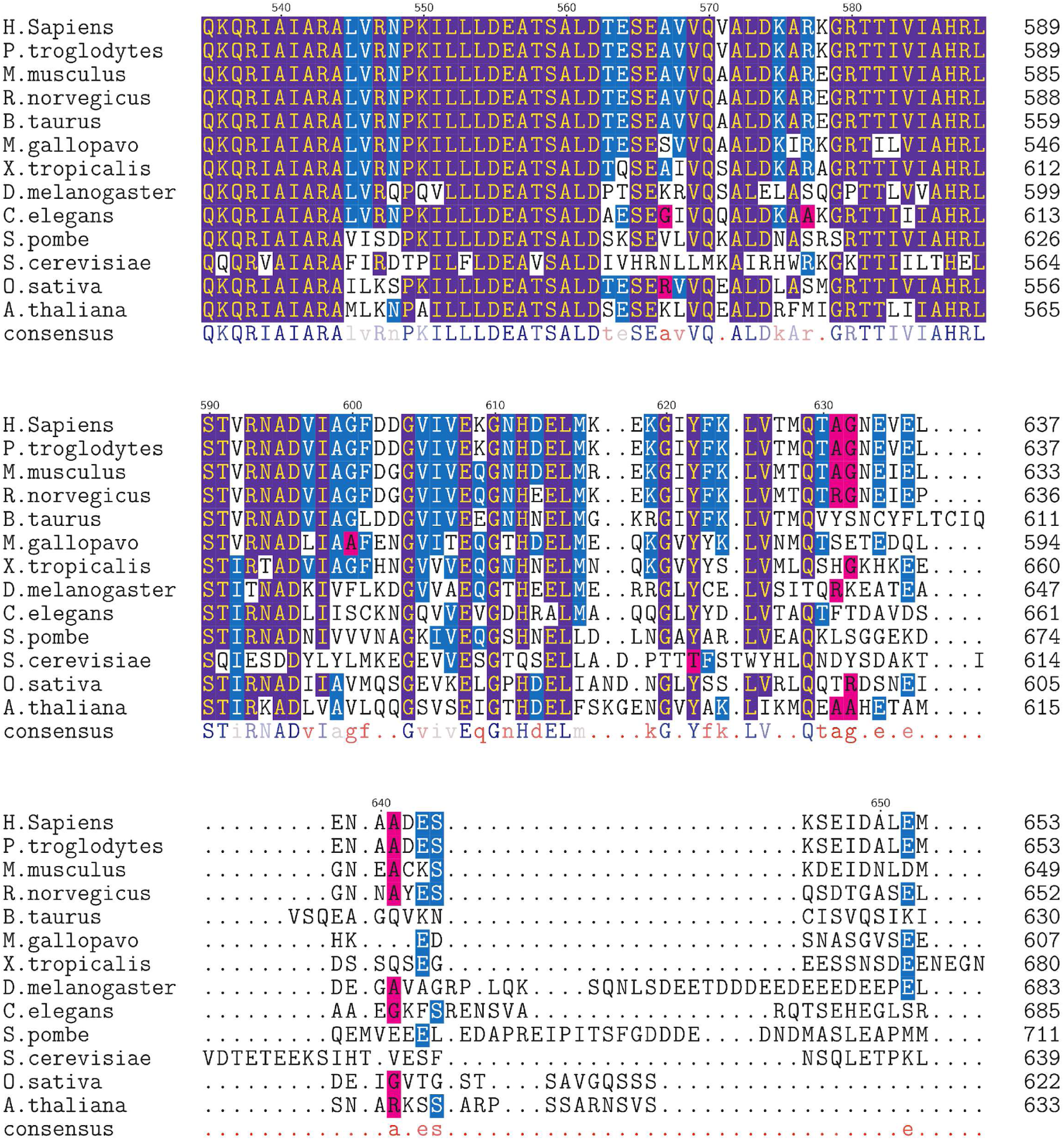
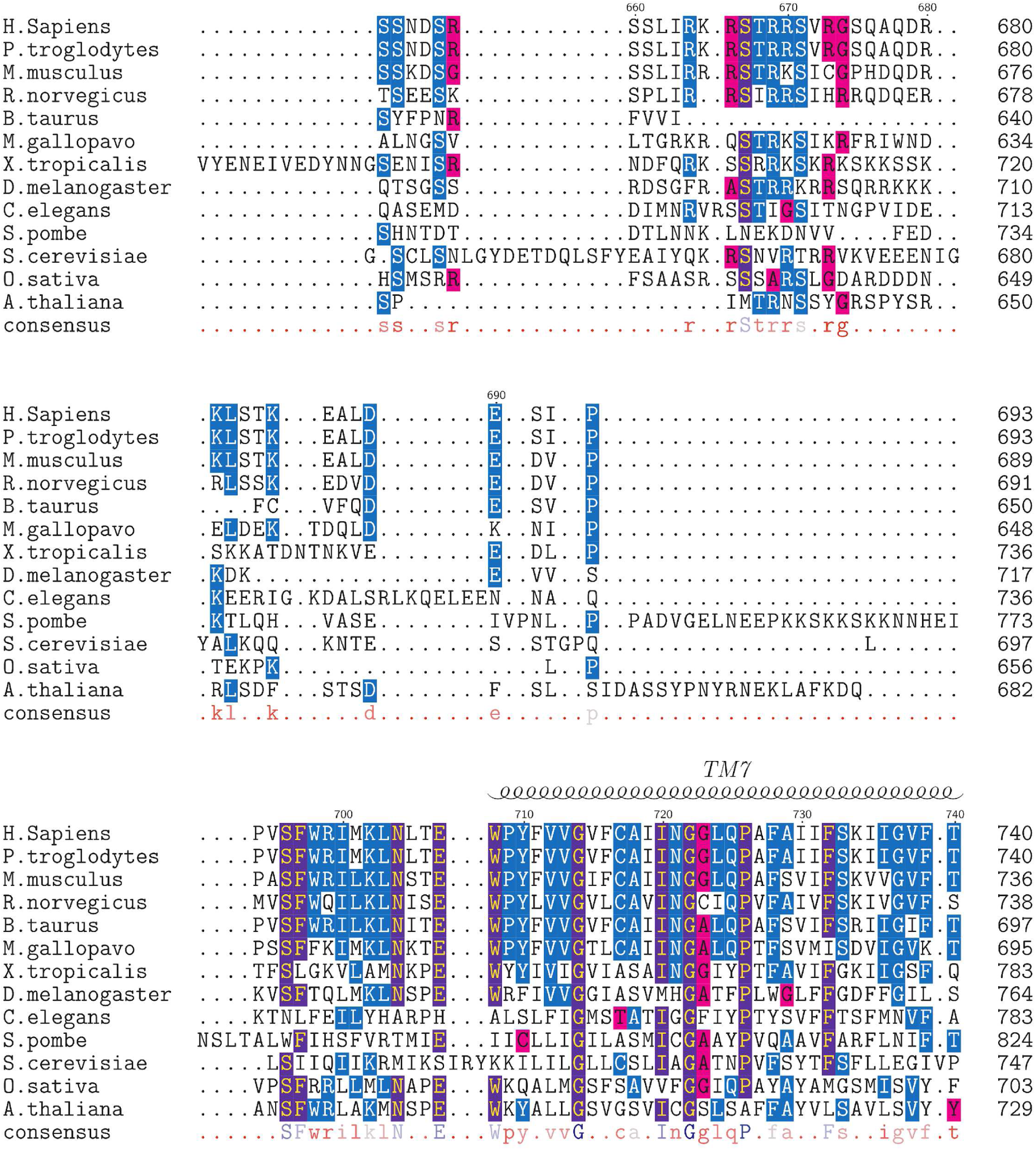
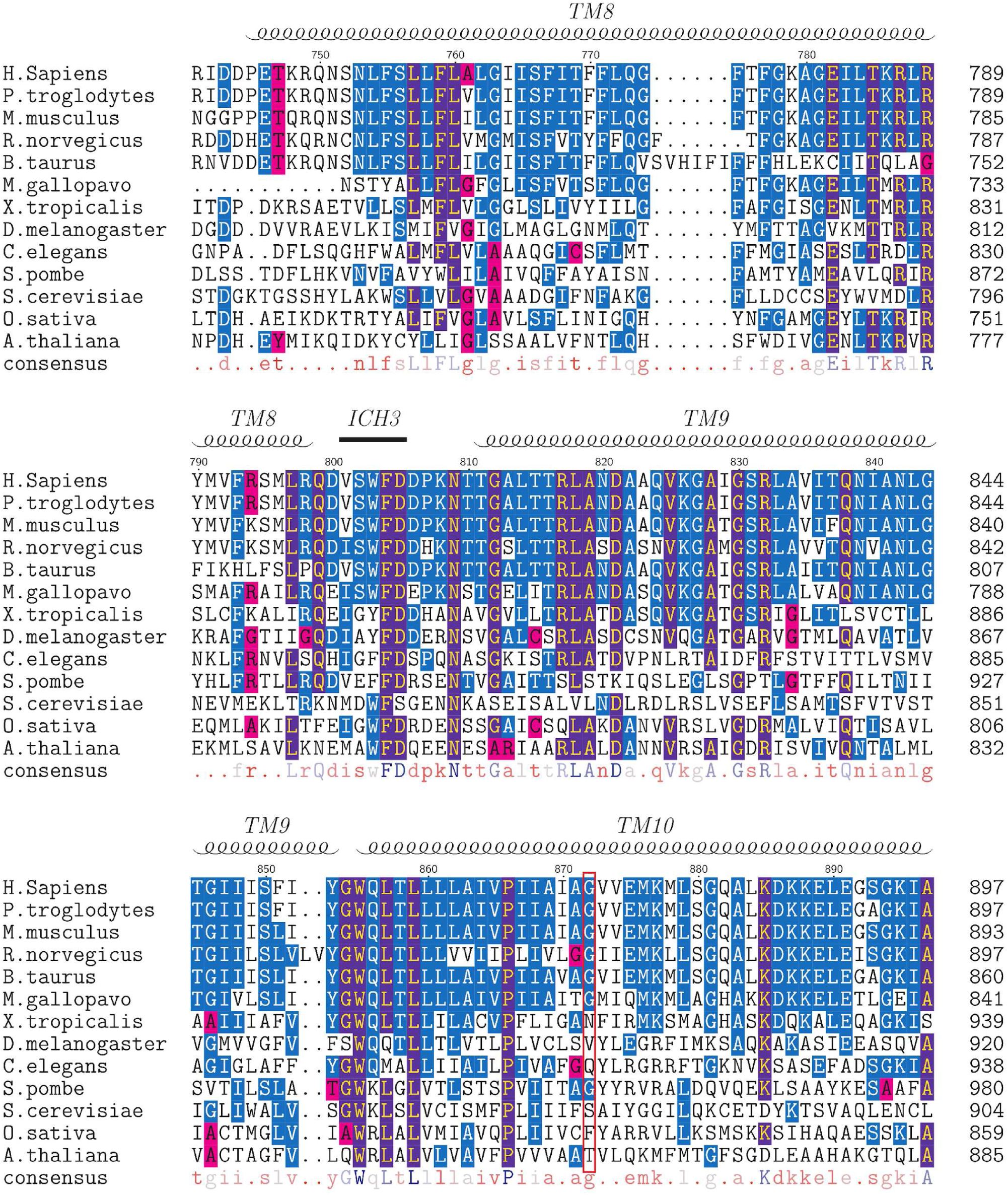
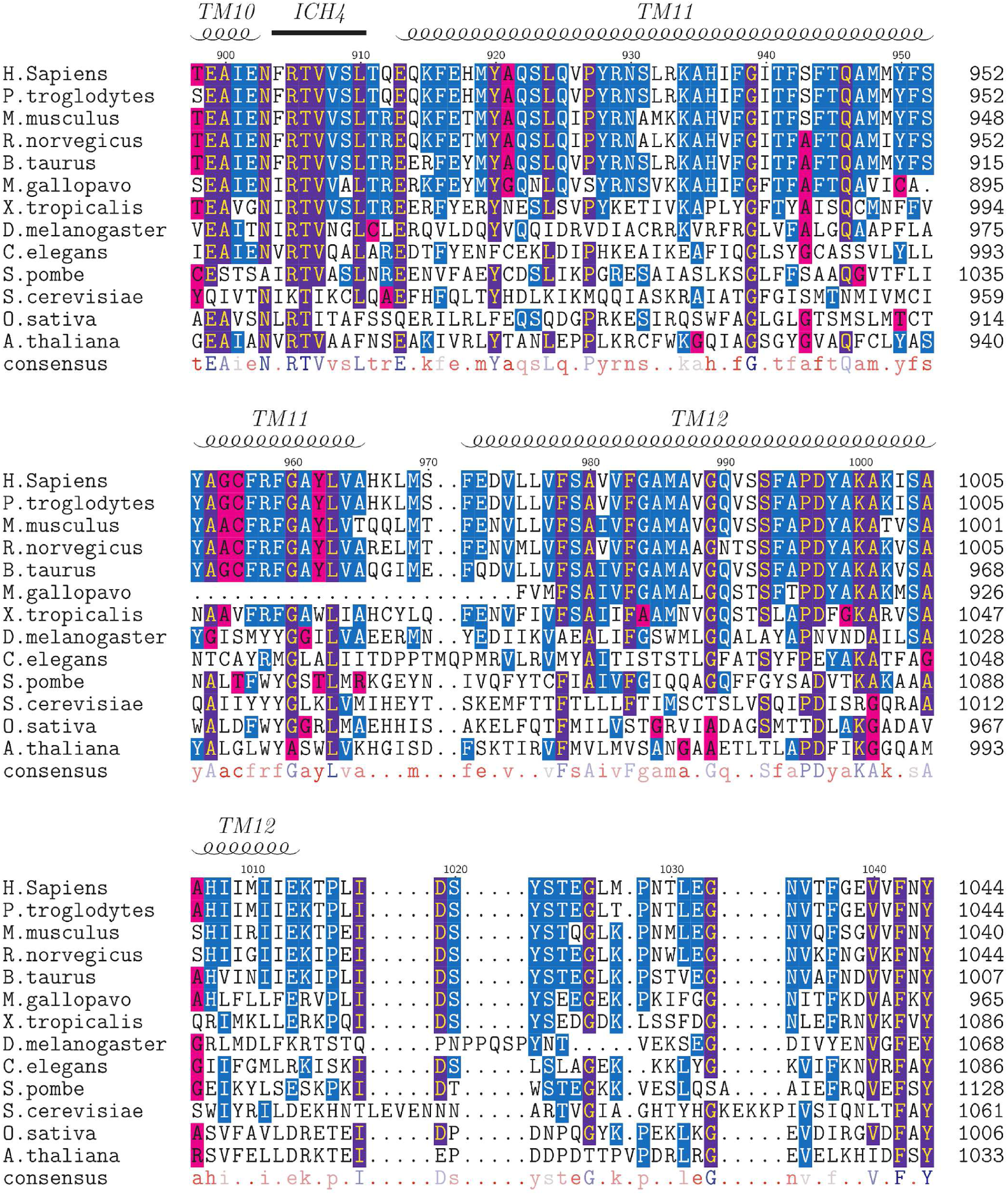
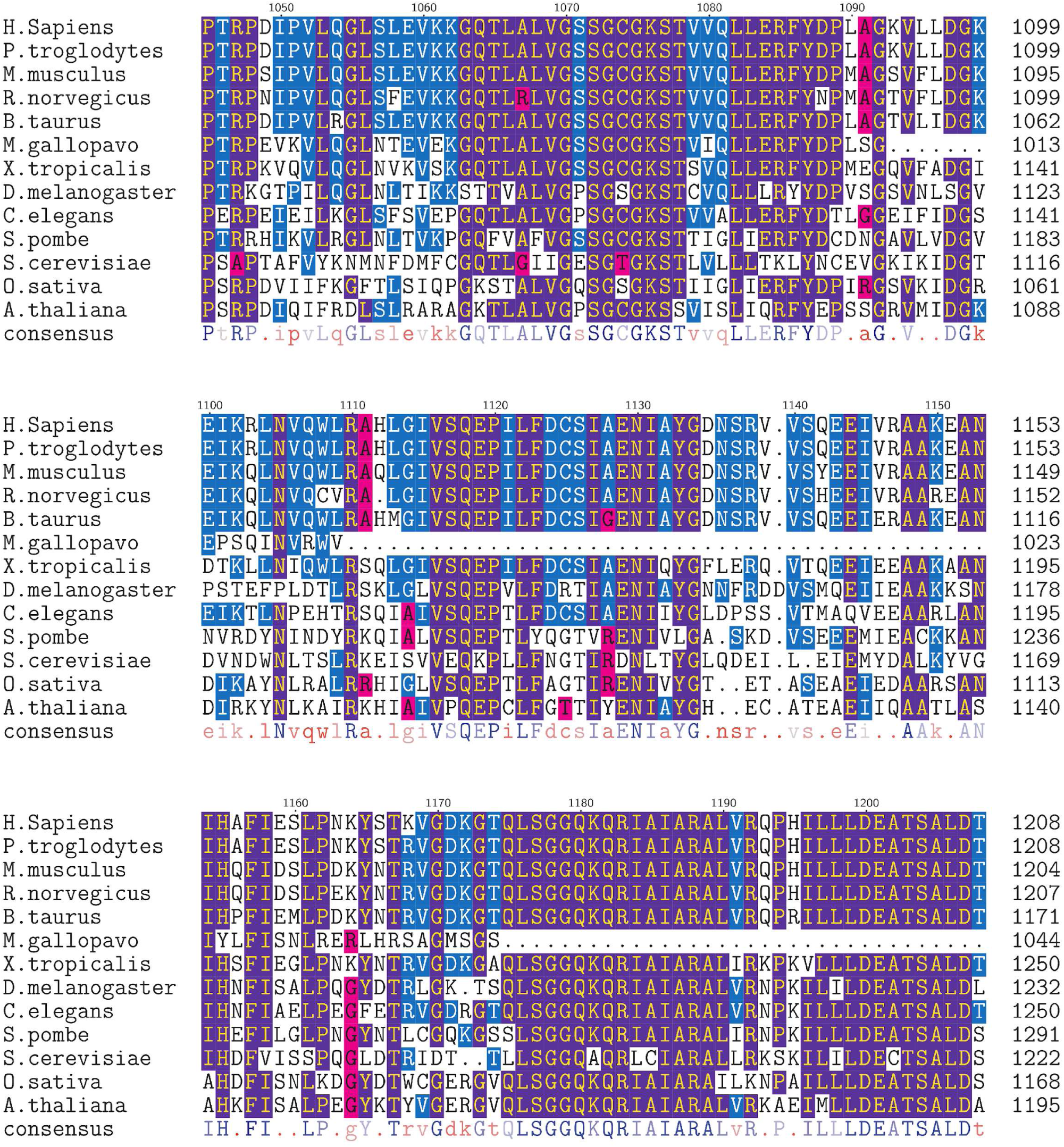
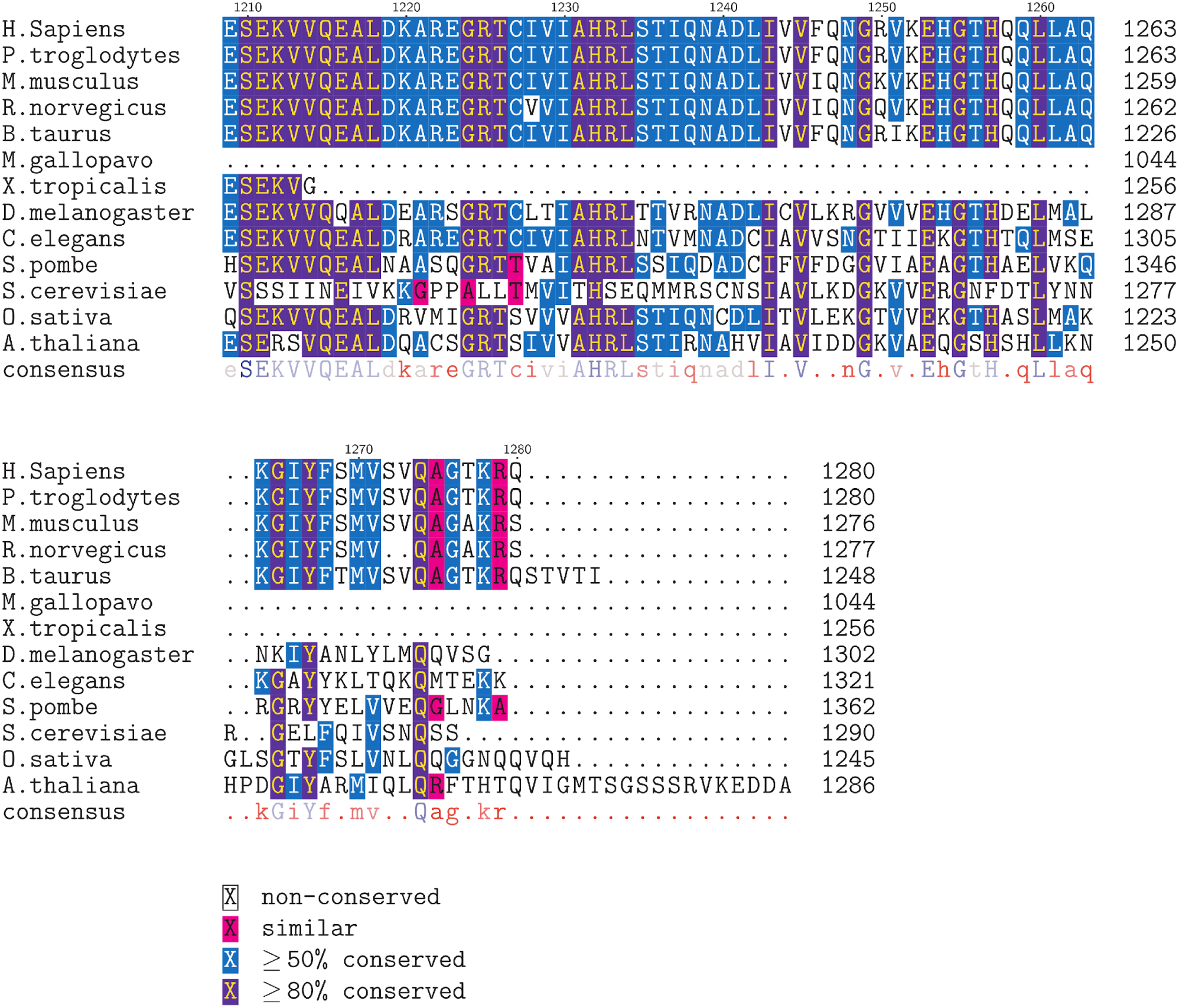
Multiple protein sequence alignment of P-gp homologous proteins. The sequence reference numbering is relative to the human P-gp. The red boxes indicate the position of the pivotal Gly226 and Gly872. The consensus is indicated below and colored from red to blue (from lower to higher degree of conservation). The color legend is indicated.

## References

(1) Vasiliou, V.; Vasiliou, K.; Nebert, D. W. Human ATP-binding Cassette (ABC) Transporter Family. Human Genomics 2009, 3, 281–290.

(2) Thomas, C.; Tampé, R. Structural and Mechanistic Principles of ABC Transporters. Annual Review of Biochemistry 2020, 89, 605–636.

(3) Orelle, C.; Schmitt, L.; Jault, J.-M. Waste or die: The price to pay to stay alive. Trends in Microbiology 2023, 31, 233–241.

(4) Alam, A.; Locher, K. P. Structure and Mechanism of Human ABC Transporters. Annual Review of Biophysics 2023, 52, 275–300.

(5) Gottesman, M. M.; Lavi, O.; Hall, M. D.; Gillet, J.-P. Toward a Better Understanding of the Complexity of Cancer Drug Resistance. Annual Review of Pharmacology and Toxicology 2016, 56, 85–102.

(6) Borst, P.; Schinkel, A. H. P-Glycoprotein ABCB1: A Major Player in Drug Handling by Mammals. The Journal of Clinical Investigation 2013, 123, 4131–4133.

(7) Eckford, P. D. W.; Sharom, F. J. The Reconstituted P-glycoprotein Multidrug Transporter Is a Flippase for Glucosylceramide and Other Simple Glycosphingolipids. Biochemical Journal 2005, 389, 517–526.

(8) Romsicki, Y.; Sharom, F. J. Phospholipid Flippase Activity of the Reconstituted Pglycoprotein Multidrug Transporter. Biochemistry 2001, 40, 6937–6947.

(9) van Helvoort, A.; Smith, A. J.; Sprong, H.; Fritzsche, I.; Schinkel, A. H.; Borst, P.; van Meer, G. MDR1 P-glycoprotein Is a Lipid Translocase of Broad Specificity, While MDR3 P-glycoprotein Specifically Translocates Phosphatidylcholine. Cell 1996, 87, 507–517.

(10) Siarheyeva, A.; Lopez, J. J.; Glaubitz, C. Localization of Multidrug Transporter Substrates within Model Membranes. Biochemistry 2006, 45, 6203–6211.

(11) Kapoor, K.; Pant, S.; Tajkhorshid, E. Active participation of membrane lipids in inhibition of multidrug transporter P-glycoprotein. Chemical Science 2021, 12, 6293–6306.

(12) Gao, Y.; Wei, C.; Luo, L.; Tang, Y.; Yu, Y.; Li, Y.; Xing, J.; Pan, X. Membrane-Assisted Tariquidar Access and Binding Mechanisms of Human ATP-binding Cassette Transporter P-glycoprotein. Frontiers in Molecular Biosciences 2024, 11, 1364494.

(13) Sharom, F. J. Complex Interplay between the P-Glycoprotein Multidrug Efflux Pump and the Membrane: Its Role in Modulating Protein Function. Frontiers in Oncology 2014, 4, 41.

(14) Peetla, C.; Vijayaraghavalu, S.; Labhasetwar, V. Biophysics of Cell Membrane Lipids in Cancer Drug Resistance: Implications for Drug Transport and Drug Delivery with Nanoparticles. Advanced drug delivery reviews 2013, 65, 10.1016/j.addr.2013.09.004.

(15) Callaghan, R.; Stafford, A.; Epand, R. M. Increased Accumulation of Drugs in a Multidrug Resistant Cell Line by Alteration of Membrane Biophysical Properties. Biochimica et Biophysica Acta (BBA) - Molecular Cell Research 1993, 1175, 277–282.

(16) Chan, L. M. S.; Lowes, S.; Hirst, B. H. The ABCs of Drug Transport in Intestine and Liver: Efflux Proteins Limiting Drug Absorption and Bioavailability. European Journal of Pharmaceutical Sciences 2004, 21, 25–51.

(17) Kroll, T.; Prescher, M.; Smits, S. H. J.; Schmitt, L. Structure and Function of Hepatobiliary ATP Binding Cassette Transporters. Chemical Reviews 2021, 121, 5240–5288.

(18) Molitoris, B. A.; Simon, F. R. Renal cortical brush-border and basolateral membranes: Cholesterol and phospholipid Composition and relative turnover. The Journal of Membrane Biology 1985, 83, 207–215.

(19) Schulze, R. J.; Schott, M. B.; Casey, C. A.; Tuma, P. L.; McNiven, M. A. The cell biology of the hepatocyte: A membrane trafficking machine. Journal of Cell Biology 2019, 218, 2096–2112.

(20) Saad, A. B.; Bruneau, A.; Mareux, E.; Lapalus, M.; Delaunay, J. L.; Gonzales, E.; Jacquemin, E.; Aït-Slimane, T.; Falguières, T. Molecular regulation of canalicular ABC transporters. International Journal of Molecular Sciences 2021, 22, 2113.

(21) Tietz, P.; Jefferson, J.; Pagano, R.; LaRusso, N. F. Membrane Microdomains in Hepatocytes: Potential Target Areas for Proteins Involved in Canalicular Bile Secretion. Journal of Lipid Research 2005, 46, 1426–1432.

(22) Kremmer, T.; Wisher, M. H.; Evans, W. H. The Lipid composition of plasma membrane subfractions originating from the three major functional domains of the rat hepatocyte cell surface. BBA - Biomembranes 1976, 455, 655–664.

(23) Modok, S.; Heyward, C.; Callaghan, R. P-Glycoprotein Retains Function When Reconstituted into a Sphingolipid- and Cholesterol-Rich Environment. Journal of Lipid Research 2004, 45, 1910–1918.

(24) Clay, A. T.; Sharom, F. J. Lipid Bilayer Properties Control Membrane Partitioning, Binding, and Transport of P-Glycoprotein Substrates. Biochemistry 2013, 52, 343–354.

(25) Romsicki, Y.; Sharom, F. J. The Membrane Lipid Environment Modulates Drug Interactions with the P-Glycoprotein Multidrug Transporter. Biochemistry 1999, 38, 6887–6896.

(26) Clay, A. T.; Lu, P.; Sharom, F. J. Interaction of the P-Glycoprotein Multidrug Transporter with Sterols. Biochemistry 2015, 54, 6586–6597.

(27) Thangapandian, S.; Kapoor, K.; Tajkhorshid, E. Probing cholesterol binding and translocation in P-glycoprotein. Biochimica et Biophysica Acta - Biomembranes 2020, 1862, 183090.

(28) Kimura, Y.; Kodan, A.; Matsuo, M.; Ueda, K. Cholesterol Fill-in Model: Mechanism for Substrate Recognition by ABC Proteins. Journal of Bioenergetics and Biomembranes 2007, 39, 447–452.

(29) Orlowski, S.; Martin, S.; Escargueil, A. P-Glycoprotein and ‘Lipid Rafts’: Some Ambiguous Mutual Relationships (Floating on Them, Building Them or Meeting Them by Chance?). Cellular and molecular life sciences: CMLS 2006, 63, 1038–1059.

(30) Kimura, Y.; Kioka, N.; Kato, H.; Matsuo, M.; Ueda, K. Modulation of Drug-Stimulated ATPase Activity of Human MDR1/P-glycoprotein by Cholesterol. Biochemical Journal 2007, 401, 597–605.

(31) Raggers, R. J.; van Helvoort, A.; Evers, R.; van Meer, G. The Human Multidrug Resistance Protein MRP1 Translocates Sphingolipid Analogs across the Plasma Membrane. Journal of Cell Science 1999, 112 *( Pt* *3*), 415–422.

(32) Veldman, R. J.; Sietsma, H.; Klappe, K.; Hoekstra, D.; Kok, J. W. Inhibition of P-glycoprotein Activity and Chemosensitization of Multidrug-Resistant Ovarian Carcinoma 2780AD Cells by Hexanoylglucosylceramide. Biochemical and Biophysical Research Communications 1999, 266, 492–496.

(33) Sietsma, H.; Veldman, R.; Kok, J. The Involvement of Sphingolipids in Multidrug Resistance. The Journal of Membrane Biology 2001, 181, 153–162.

(34) Cannon, R. E.; Peart, J. C.; Hawkins, B. T.; Campos, C. R.; Miller, D. S. Targeting Blood-Brain Barrier Sphingolipid Signaling Reduces Basal P-glycoprotein Activity and Improves Drug Delivery to the Brain. Proceedings of the National Academy of Sciences of the United States of America 2012, 109, 15930–15935.

(35) Alam, A.; Küng, R.; Kowal, J.; McLeod, R. A.; Tremp, N.; Broude, E. V.; Roninson, I. B.; Stahlberg, H.; Locher, K. P. Structure of a zosuquidar and UIC2-bound human-mouse chimeric ABCB1. Proceedings of the National Academy of Sciences of the United States of America 2018, 115, E1973–E1982.

(36) Gewering, T.; Waghray, D.; Parey, K.; Jung, H.; Tran, N. N.; Zapata, J.; Zhao, P.; Chen, H.; Januliene, D.; Hummer, G.; Urbatsch, I.; Moeller, A.; Zhang, Q. Tracing the Substrate Translocation Mechanism in P-glycoprotein. eLife 2024, 12, RP90174.

(37) Barbieri, A.; Thonghin, N.; Shafi, T.; Prince, S. M.; Collins, R. F.; Ford, R. C. Structure of ABCB1/P-Glycoprotein in the Presence of the CFTR Potentiator Ivacaftor. Membranes 2021, 11, 923.

(38) Le, C. A.; Harvey, D. S.; Aller, S. G. Structural Definition of Polyspecific Compensatory Ligand Recognition by P-glycoprotein. IUCrJ 2020, 7, 663–672.

(39) Thonghin, N.; Collins, R. F.; Barbieri, A.; Shafi, T.; Siebert, A.; Ford, R. C. Novel Features in the Structure of P-glycoprotein (ABCB1) in the Post-Hydrolytic State as Determined at 7.9 Å Resolution. BMC structural biology 2018, 18, 17.

(40) Esser, L.; Zhou, F.; Pluchino, K. M.; Shiloach, J.; Ma, J.; Tang, W.-K.; Gutierrez, C.; Zhang, A.; Shukla, S.; Madigan, J. P.; Zhou, T.; Kwong, P. D.; Ambudkar, S. V.; Gottesman, M. M.; Xia, D. Structures of the Multidrug Transporter P-glycoprotein Reveal Asymmetric ATP Binding and the Mechanism of Polyspecificity. The Journal of Biological Chemistry 2017, 292, 446–461.

(41) Nicklisch, S. C. T.; Rees, S. D.; McGrath, A. P.; Gökirmak, T.; Bonito, L. T.; Vermeer, L. M.; Cregger, C.; Loewen, G.; Sandin, S.; Chang, G.; Hamdoun, A. Global Marine Pollutants Inhibit P-glycoprotein: Environmental Levels, Inhibitory Effects, and Cocrystal Structure. Science Advances 2016, 2, e1600001.

(42) Szewczyk, P.; Tao, H.; McGrath, A. P.; Villaluz, M.; Rees, S. D.; Lee, S. C.; Doshi, R.; Urbatsch, I. L.; Zhang, Q.; Chang, G. Snapshots of Ligand Entry, Malleable Binding and Induced Helical Movement in P-glycoprotein. Acta Crystallographica. Section D, Biological Crystallography 2015, 71, 732–741.

(43) Kodan, A.; Yamaguchi, T.; Nakatsu, T.; Matsuoka, K.; Kimura, Y.; Ueda, K.; Kato, H. Inward-and Outward-Facing X-ray Crystal Structures of Homodimeric P-glycoprotein CmABCB1. Nature Communications 2019, 10, 88.

(44) Kim, Y.; Chen, J. Molecular structure of human P-glycoprotein in the ATP-bound, outward-facing conformation. Science 2018, 359, 915–919.

(45) Jin, M. S.; Oldham, M. L.; Zhang, Q.; Chen, J. Crystal Structure of the Multidrug Transporter P-glycoprotein from Caenorhabditis Elegans. Nature 2012, 490, 566–569.

(46) Ward, A.; Reyes, C. L.; Yu, J.; Roth, C. B.; Chang, G. Flexibility in the ABC Transporter MsbA: Alternating Access with a Twist. Proceedings of the National Academy of Sciences of the United States of America 2007, 104, 19005–19010.

(47) Galazzo, L.; Meier, G.; Januliene, D.; Parey, K.; De Vecchis, D.; Striednig, B.; Hilbi, H.; Schäfer, L. V.; Kuprov, I.; Moeller, A.; Bordignon, E.; Seeger, M. A. The ABC transporter MsbA adopts the wide inward-open conformation in E. coli cells. Science Advances 2022, 8, eabn6845.

(48) Nosol, K.; Romane, K.; Irobalieva, R. N.; Alam, A.; Kowal, J.; Fujita, N.; Locher, K. P. Cryo-EM structures reveal distinct mechanisms of inhibition of the human multidrug transporter ABCB1. Proceedings of the National Academy of Sciences 2020, 117, 26245–26253.

(49) Urgaonkar, S.; Nosol, K.; Said, A. M.; Nasief, N. N.; Bu, Y.; Locher, K. P.; Lau, J. Y. N.; Smolinski, M. P. Discovery and Characterization of Potent Dual P-Glycoprotein and CYP3A4 Inhibitors: Design, Synthesis, Cryo-EM Analysis, and Biological Evaluations. Journal of Medicinal Chemistry 2022, 65, 191–216.

(50) Alam, A.; Kowal, J.; Broude, E.; Roninson, I.; Locher, K. P. Structural insight into substrate and inhibitor discrimination by human P-glycoprotein. Science 2019, 363, 753–756.

(51) Ellinghaus, T. L.; Marcellino, T.; Srinivasan, V.; Lill, R.; Kühlbrandt, W. Conformational Changes in the Yeast Mitochondrial ABC Transporter Atm1 during the Transport Cycle. Science Advances 2021, 7, eabk2392.

(52) Lindahl, E.; Sansom, M. S. P. Membrane proteins: molecular dynamics simulations. Current Opinion in Structural Biology 2008, 18, 425–431.

(53) Li, J.; Wen, P.-C.; Moradi, M.; Tajkhorshid, E. Computational characterization of structural dynamics underlying function in active membrane transporters. Current Opinion in Structural Biology 2015, 31, 96–105.

(54) Szöllösi, D.; Rose-Sperling, D.; Hellmich, U. A.; Stockner, T. Comparison of mechanistic transport cycle models of ABC exporters. Biochimica et Biophysica Acta (BBA) - Biomembranes 2018, 1860, 818–832.

(55) Furuta, T. Structural dynamics of ABC transporters: molecular simulation studies. Biochemical Society Transactions 2021, 49, 405–414.

(56) Badiee, S. A.; Isu, U. H.; Khodadadi, E.; Moradi, M. The Alternating Access Mechanism in Mammalian Multidrug Resistance Transporters and Their Bacterial Homologs. Membranes 2023, 13, 568.

(57) Göddeke, H.; Timachi, M. H.; Hutter, C. A.; Galazzo, L.; Seeger, M. A.; Karttunen, M.; Bordignon, E.; Schäfer, L. V. Atomistic Mechanism of Large-Scale Conformational Transition in a Heterodimeric ABC Exporter. Journal of the American Chemical Society 2018, 140, 4543–4551.

(58) Göddeke, H.; Schäfer, L. V. Capturing Substrate Translocation in an ABC Exporter at the Atomic Level. Journal of the American Chemical Society 2020, 142, 12791–12801.

(59) Wen, P.-C.; Verhalen, B.; Wilkens, S.; Mchaourab, H. S.; Tajkhorshid, E. On the Origin of Large Flexibility or P-glycoprotein in the Inward-facing State. Journal of Biological Chemistry 2013, 228, 19211–19220.

(60) Barreto-Ojeda, E.; Corradi, V.; Gu, R. X.; Tieleman, D. P. Coarse-grained molecular dynamics simulations reveal lipid access pathways in P-glycoprotein. Journal of General Physiology 2018, 150, 417–429.

(61) Domicevica, L.; Koldsø, H.; Biggin, P. C. Multiscale molecular dynamics simulations of lipid interactions with P-glycoprotein in a complex membrane. Journal of Molecular Graphics and Modelling 2018, 80, 147–156.

(62) McCormick, J. W.; Vogel, P. D.; Wise, J. G. Multiple Drug Transport Pathways through Human P-Glycoprotein. Biochemistry 2015, 54, 4374–4390.

(63) Xing, J.; Mei, H.; Huang, S.; Zhang, D.; Pan, X. An Energetically Favorable Ligand Entrance Gate of a Multidrug Transporter Revealed by Partial Nudged Elastic Band Simulations. Computational and Structural Biotechnology Journal 2019, 17, 319–323.

(64) Nasim, F.; Schmid, D.; Szakács, G.; Sohail, A.; Sitte, H. H.; Chiba, P.; Stockner, T. Active Transport of Rhodamine 123 by the Human Multidrug Transporter P-glycoprotein Involves Two Independent Outer Gates. Pharmacology Research & Perspectives 2020, 8, e00572.

(65) Mora Lagares, L.; Pérez-Castillo, Y.; Minovski, N.; Novic, M. Structure–Function Relationships in the Human P-Glycoprotein (ABCB1): Insights from Molecular Dynamics Simulations. International Journal of Molecular Sciences 2022, 23, 362.

(66) Jorgensen, C.; Ulmschneider, M. B.; Searson, P. C. Modeling Substrate Entry into the P-Glycoprotein Efflux Pump at the Blood-Brain Barrier. Journal of Medicinal Chemistry 2023, 66, 16615–16627.

(67) Focht, D.; Croll, T. I.; Pedersen, B. P.; Nissen, P. Improved Model of Proton Pump Crystal Structure Obtained by Interactive Molecular Dynamics Flexible Fitting Expands the Mechanistic Model for Proton Translocation in P-Type ATPases. Frontiers in Physiology 2017, 8, 202.

(68) Williams, C. J. et al. MolProbity: More and better reference data for improved all-atom structure validation. Protein Science 2018, 27, 293–315.

(69) Webb, B.; Sali, A. Comparative protein structure modeling using MODELLER. Current Protocols in Bioinformatics 2016, 2016, 5.6.1–5.6.37.

(70) Li, J.; Jaimes, K. F.; Aller, S. G. Refined structures of mouse P-glycoprotein. Protein Science 2014, 23, 34–46.

(71) Abraham, M. J.; Murtola, T.; Schulz, R.; Páll, S.; Smith, J. C.; Hess, B.; Lindahl, E. GROMACS: High performance molecular simulations through multi-level parallelism from laptops to supercomputers. SoftwareX 2015, 1–2, 19–25.

(72) De Jong, D. H.; Singh, G.; Bennett, W. F.; Arnarez, C.; Wassenaar, T. A.; Scháfer, L. V.; Periole, X.; Tieleman, D. P.; Marrink, S. J. Improved parameters for the martini coarse-grained protein force field. Journal of Chemical Theory and Computation 2013, 9, 687–697.

(73) Bussi, G.; Donadio, D.; Parrinello, M. Canonical sampling through velocity rescaling. The Journal of Chemical Physics 2007, 126, 014101.

(74) Bernetti, M.; Bussi, G. Pressure control using stochastic cell rescaling. The Journal of Chemical Physics 2020, 153, 114107.

(75) Wassenaar, T. A.; Pluhackova, K.; Böckmann, R. A.; Marrink, S. J.; Tieleman, D. P. Going Backward: A Flexible Geometric Approach to Reverse Transformation from Coarse Grained to Atomistic Models. Journal of Chemical Theory and Computation 2014, 10, 676–690.

(76) Huang, J.; Rauscher, S.; Nawrocki, G.; Ran, T.; Feig, M.; De Groot, B. L.; Grubmüller, H.; MacKerell, A. D. CHARMM36m: An improved force field for folded and intrinsically disordered proteins. Nature Methods 2016, 14, 71–73.

(77) Allnér, O.; Nilsson, L.; Villa, A. Magnesium ion-water coordination and exchange in biomolecular simulations. Journal of Chemical Theory and Computation 2012, 8, 1493– 1502.

(78) Darden, T.; York, D.; Pedersen, L. Particle mesh Ewald: An N log(N) method for Ewald sums in large systems. The Journal of Chemical Physics 1993, 98, 10089–10092.

(79) Hess, B.; Bekker, H.; Berendsen, H. J. C.; Fraaije, J. G. E. M. LINCS: A linear constraint solver for molecular simulations. Journal of Computational Chemistry 1997, 18, 1463–1472.

(80) Hess, B. P-LINCS: A parallel linear constraint solver for molecular simulation. Journal of Chemical Theory and Computation 2008, 4, 116–122.

(81) Notredame, C.; Higgins, D. G.; Heringa, J. T-coffee: a novel method for fast and accurate multiple sequence alignment 1 1Edited by J. Thornton. Journal of Molecular Biology 2000, 302, 205–217.

(82) Consortium, T. U. UniProt: A Hub for Protein Information. Nucleic Acids Research 2015, 43, D204–D212.

(83) Tate, J. G. et al. COSMIC: The Catalogue Of Somatic Mutations In Cancer. Nucleic Acids Research 2019, 47, D941–D947.

(84) Paramo, T.; East, A.; Garzón, D.; Ulmschneider, M. B.; Bond, P. J. Efficient Characterization of Protein Cavities within Molecular Simulation Trajectories: trj cavity. Journal of Chemical Theory and Computation 2014, 10, 2151–2164.

(85) Joosten, R. P.; Te Beek, T. A.; Krieger, E.; Hekkelman, M. L.; Hooft, R. W.; Schneider, R.; Sander, C.; Vriend, G. A series of PDB related databases for everyday needs. Nucleic Acids Research 2011, 39, D411–D419.

(86) Corradi, V.; Mendez-Villuendas, E.; Ingólfsson, H. I.; Gu, R.-X.; Siuda, I.; Melo, M. N.; Moussatova, A.; DeGagné, L. J.; Sejdiu, B. I.; Singh, G.; Wassenaar, T. A.; Delgado Magnero, K.; Marrink, S. J.; Tieleman, D. P. Lipid–Protein Interactions Are Unique Fingerprints for Membrane Proteins. ACS Central Science 2018, 4, 709–717.

(87) Loo, T. W.; Clarke, D. M. Mapping the Binding Site of the Inhibitor Tariquidar That Stabilizes the First Transmembrane Domain of P-glycoprotein. Journal of Biological Chemistry 2015, 290, 29389–29401.

(88) Elferink, R. P.; Tytgat, G. N.; Groen, A. K. Hepatic Canalicular Membrane 1: The Role of Mdr2 P-glycoprotein in Hepatobiliary Lipid Transport. FASEB Journal 1997, 11, 19–28.

(89) Alam, A.; Kowal, J.; Broude, E.; Roninson, I.; Locher, K. P. Structural Insight into Substrate and Inhibitor Discrimination by Human P-glycoprotein. *Science (New York*, N.Y.) 2019, 363, 753–756.

(90) Wang, L.; O’Mara, M. L. Effect of the Force Field on Molecular Dynamics Simulations of the Multidrug Efflux Protein P-Glycoprotein. Journal of Chemical Theory and Computation 2021, 17, 6491–6508.

(91) Dastvan, R.; Mishra, S.; Peskova, Y. B.; Nakamoto, R. K.; Mchaourab, H. S. Mechanism of allosteric modulation of P-glycoprotein by transport substrates and inhibitors. Science 2019, 364, 689–692.

(92) Verhalen, B.; Dastvan, R.; Thangapandian, S.; Peskova, Y.; Koteiche, H. A.; Nakamoto, R. K.; Tajkhorshid, E.; McHaourab, H. S. Energy transduction and alternating access of the mammalian ABC transporter P-glycoprotein. Nature 2017, 543, 738–741.

(93) Debruycker, V.; Hutchin, A.; Masureel, M.; Ficici, E.; Martens, C.; Legrand, P.; Stein, R.; Mchaourab, H.; Faraldo-Gómez, J. D.; Remaut, H.; Govaerts, C. An Embedded Lipid in the Multidrug Transporter LmrP Suggests a Mechanism for Polyspecificity. Nature structural & molecular biology 2020, 27, 829–835.

